# HPV-Associated Tonsillar Intraepithelial Neoplasia and Oral High-Risk HPV mRNA as a Candidate Biomarker

**DOI:** 10.64898/2026.10.16.759752

**Authors:** Masashi Mori, Hidenori Tanaka, Masami Suzuki, Kansuke Kido, Kazuya Ogawa, Noah Sasa, Toshihiro Kishikawa, Nagayasu Egawa, Daichi Maeda, Masafumi Horie, Kiyohito Hosokawa, Motoyuki Suzuki, Hiroshi Kato, Kazuki Hayashi, Syusuke Okamoto, Tadashi Yoshii, Shotaro Harada, Tetsuo Morihana, Sho Obata, Satoru Uetsuka, Suetaka Nishiike, Naoki Ashida, Rumi Kambara, Tsutomu Sano, Kayoko Higashi-Shingai, Takayuki Kawashima, Aya Kamakura, Kenji Mitani, Hayato Mori, Rentaro Watanabe, Tomoki Yasuoka, Eiichi Morii, Osaka Study Consortium for Tonsillar Intraepithelial Neoplasia, Hidenori Inohara

## Abstract

Early detection of human papillomavirus (HPV)-associated oropharyngeal cancer is an unmet need, as its premalignant lesions remain unestablished. We hypothesized that such lesions exist in individuals with oral high-risk HPV (HR-HPV) mRNA. We preoperatively screened 1,180 individuals undergoing tonsillectomy due to benign conditions and detected five with oral HR-HPV mRNA. Serial sectioning of the entire excised tonsils identified HPV-associated intraepithelial lesions in three of these five individuals. A similar lesion was identified in an additional individual with oral HPV16 infection with unknown oral HR-HPV mRNA status. Intraepithelial lesions in two individuals were HPV16-associated and showed moderate-to-severe atypia, whereas those in the other two individuals were associated with other HR-HPV and showed minimal-to-mild atypia. Based on HPV life cycle, the former were classified as transforming lesions and the latter as productive-to-intermediate lesions. Spatial transcriptomic analysis revealed molecular profiles intermediate between those of normal epithelium and HPV-associated oropharyngeal cancer. These findings establish the first evidence for HPV-associated tonsillar premalignant lesions and suggest oral HR-HPV mRNA as a candidate biomarker.

## INTRODUCTION

Human papillomavirus (HPV)-associated cancers, such as cervical and oropharyngeal cancers, account for 5% of all malignancies worldwide.^1,2^ HPV-associated cancers can be eradicated through primary prevention by HPV vaccination and secondary prevention by early detection. However, HPV-associated oropharyngeal cancer is increasing rapidly, particularly in developed countries, and its incidence has already surpassed that of cervical cancer in the United States and the United Kingdom.^3,4^ A similar trend is observed in European countries, including Germany, Sweden, and Denmark.^5–7^ This likely reflects a greater difficulty in early detection of HPV-associated oropharyngeal cancer in contrast to cervical cancer.^8^

Direct visualization and sampling of the cervix have defined the natural history of HPV infection and enabled understanding of the carcinogenic process, from HPV infection through premalignant lesions to invasive cancer.^9,10^ Methods for early detection of cervical cancer have been well established and screening is conducted around the world, primarily in high-income countries.^11–14^ Premalignant lesions of the cervix are designated as cervical intraepithelial neoplasia (CIN) and graded as CIN 1, 2, and 3 according to the degree of dysplasia.^9^ In contrast, HPV-associated oropharyngeal cancer arises from the tonsillar crypts,^15^ which are difficult to directly visualize and sample.^16^ The natural history of HPV infection in the oropharynx remains unclear, and precursors of HPV-associated oropharyngeal cancer have never been demonstrated.^8,17^ Achieving early detection of this cancer, therefore, requires establishing the existence of premalignant lesions and identifying biomarkers for their detection.^3^

The epithelium of the tonsillar crypts is reticular and has a discontinuous basement membrane, allowing intraepithelial lesions to access lymphatics without conventional stromal invasion.^18–20^ In patients with cancer of unknown primary, occult HPV-associated tonsillar lesions are often confined to the epithelium despite neck nodal metastases.^21^ Given these findings, any HPV-associated tonsillar neoplasm is currently considered potentially malignant regardless of invasion beyond the basement membrane.^22,23^ Premalignancy therefore requires not only histological dysplasia but also molecular biological evidence that the lesion is distinct from cancer.^24^

The most definitive method for confirming HPV-associated premalignant oropharyngeal lesions is to thoroughly examine tonsils from cancer-free individuals with oral high-risk HPV (HR-HPV) infection. Oral HPV infection is assessed by detecting HPV DNA in oral rinses or gargles. However, most oral HPV infections are transient,^25^ and HPV DNA alone cannot distinguish transient, clinically irrelevant infection from persistent, active infection with malignant potential. Screening individuals solely on the basis of oral HR-HPV infection is therefore insufficient. HR-HPV E6/E7 oncoproteins are essential for HPV-driven carcinogenesis,^26,27^ and cervical HR-HPV E6/E7 mRNA levels increase as carcinogenesis progresses; HR-HPV E6/E7 mRNA is used as a biomarker of cervical neoplasia.^28,29^ Oral HR-HPV E6/E7 mRNA is detected in patients with HPV-associated oropharyngeal cancer^30^ and in some cancer-free individuals with oral HR-HPV infection,^31,32^ but not in those without infection. Individuals with oral HR-HPV E6/E7 mRNA may therefore be at increased risk of HPV-associated oropharyngeal cancer. We hypothesized that HPV-associated premalignant lesions are present in the tonsils of cancer-free individuals with oral HR-HPV E6/E7 mRNA. To identify such lesions, we decided to prepare serial sections of the entire excised tonsils. However, because this process is extremely labor-intensive, we focused on individuals with oral HR-HPV infection who tested positive for oral HR-HPV E6/E7 mRNA.

## RESULTS

### Oral HR-HPV DNA and mRNA and ctHPVDNA

Fig. 1 illustrates the workflow for the identification of latent HPV-associated tonsillar lesions. Overall, 1,180 cancer-free individuals who underwent tonsillectomy for benign conditions between February 2018 and March 2022 were screened preoperatively for oral HR-HPV. Baseline characteristics are shown in Extended Data Table 1. The median age was 30 years, and the male-to-female ratio was 1:1. Recurrent tonsillitis was the most common indication for tonsillectomy (81%), followed by IgA nephropathy (14%). The median interval between the first and second gargle collection was 31 days.

**Fig. 1:**
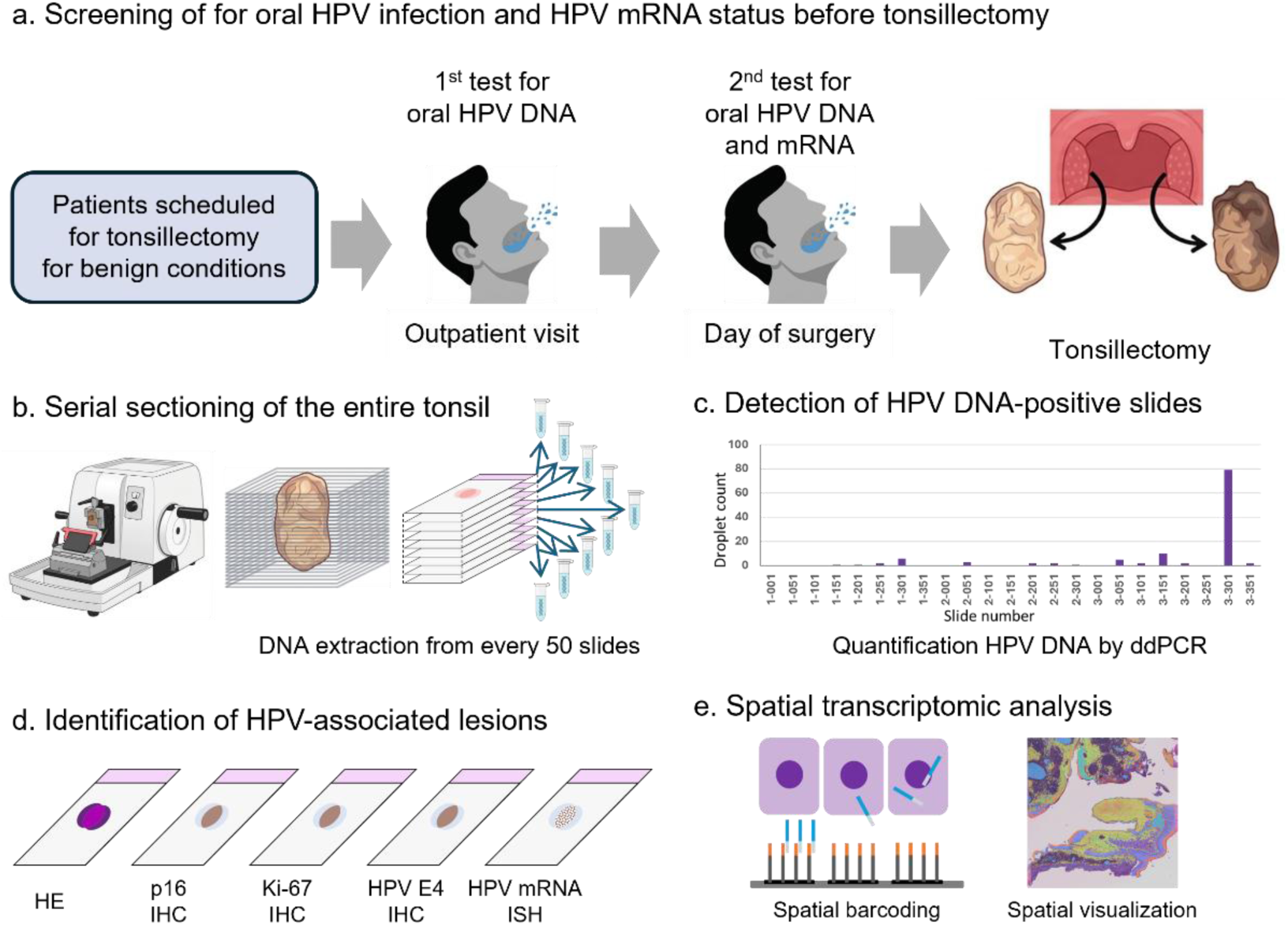
Workflow for identifying HPV-associated lesions in the tonsil. **a.** Patients scheduled for tonsillectomy for benign conditions were tested for oral HR-HPV DNA before surgery (first test); those with a positive result underwent a second test for oral HR-HPV DNA and mRNA immediately before surgery. **b.** Excised tonsils were serially sectioned, and DNA was extracted at fixed intervals. **c.** Droplet digital PCR identified slides containing HR-HPV DNA matching the oral HPV genotype. **d.** Adjacent slides were examined using hematoxylin and eosin (HE) staining, immunohistochemistry (IHC) for p16, Ki-67, and HPV E4, and *in situ* hybridization (ISH) for HR-HPV E6/E7 mRNA. **e.** Identified lesions underwent spatial transcriptomic analysis. HPV, human papillomavirus; HR-HPV, high-risk HPV.

Fig. 2 shows a flowchart of the oral HPV test results. In the first test, 28 of the 1,180 patients tested positive for oral HR-HPV DNA. The prevalence of oral HR-HPV infection was 2.4% (95% confidence interval [CI], 1.6–3.4), including 0.3% of HPV16 (95% CI, 0.1-0.7) and 2.1% other HR-HPV types (95% CI, 1.4-3.1). Compared with patients in their 20s, oral HR-HPV infection was more prevalent in those in their 40s (5.0% vs. 1.1%; odds ratio, 4.9; 95% CI, 1.7–14.1; P<0.01) and 50s (7.9% vs. 1.1%; odds ratio, 8.0; 95% CI, 2.4–27.1; P<0.01). Prevalence was higher in patients with recurrent tonsillitis than in those with other non-infectious conditions (2.8% vs. 0.5%; odds ratio, 6.4; 95% CI, 1.0–263.3; P<0.05). Prevalence was also higher in men than in women, although not significantly (3.2% vs. 1.6%; odds ratio, 2.1; 95% CI, 0.9–4.6; P=0.09) (Extended Data Table 2).

**Fig. 2:**
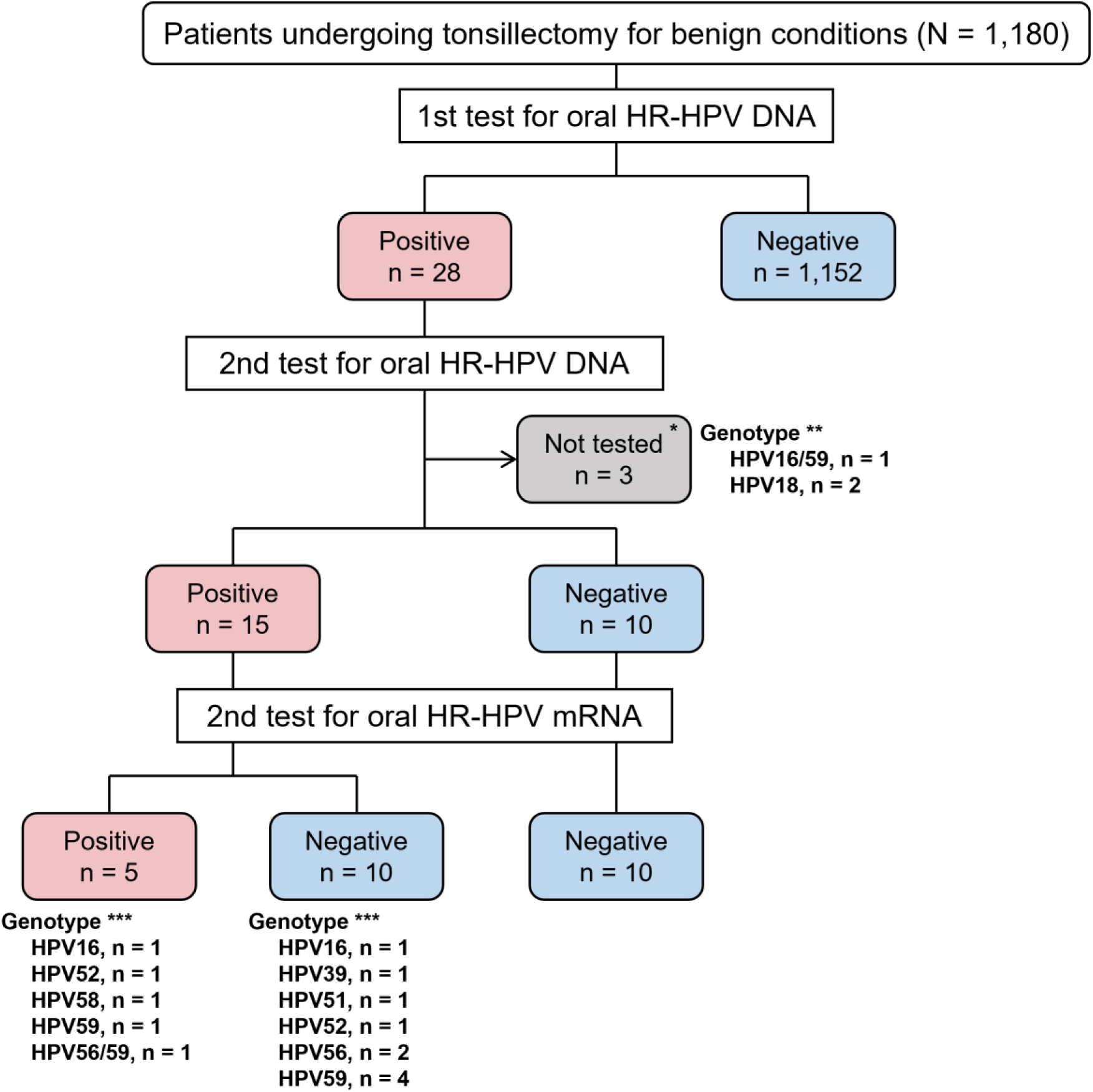
Flowchart of oral high-risk HPV DNA and mRNA test results. *Three patients did not undergo the second oral HPV DNA/mRNA test: tonsillectomy was canceled in one, whereas two underwent tonsillectomy without collection of a second gargle specimen. **HR-HPV genotype detected in the first test. ***HR-HPV genotype detected in the second test. HPV, human papillomavirus; HR-HPV, high-risk HPV.

The second test assessed persistent oral HR-HPV infection and oral HR-HPV mRNA status. Of the 28 patients positive in the first test, 25 underwent the second test. Of the three patients who did not undergo the second test, one had tonsillectomy canceled, and two underwent tonsillectomy without collection of a second gargle specimen. Fifteen of 25 patients (60.0%) remained positive for oral HR-HPV DNA, and five of these 15 (33.3%) were positive for oral HR-HPV mRNA. All patients negative for oral HR-HPV DNA on the second test were also negative for HR-HPV mRNA; thus, oral HR-HPV mRNA was detected only with persistent infection. Persistent infection was more frequent for HPV16 (100%; 95% CI, 15.8–100) than for other HR-HPV types (56.5%; 95% CI, 34.5–76.8), although the difference was not significant (P=0.50). Among patients with persistent infection, HR-HPV mRNA detection was higher for HPV16 (50.0%; 95% CI, 1.3–98.7) than for other HR-HPV types (30.8%; 95% CI, 9.1–61.4), but the difference was not significant (P=1.00) (Extended Data Table 3). Oral HPV DNA genotypes and HR-HPV mRNA status for each patient are listed in Extended Data Table 4. All HPV DNA genotypes detected in the second test had also been detected in the first. All patients tested for circulating tumor HPV DNA (ctHPVDNA) were negative.

### HR-HPV DNA-positive sites in the tonsil

We prepared serial sections of the entire tonsil excised from patients positive for oral HR-HPV mRNA and from patients with oral HPV16 infection regardless of mRNA status because HPV16 causes most HPV-associated oropharyngeal cancers.^33^ Seven patients met these criteria: Patient 1, HPV16 DNA-positive/mRNA-positive; Patient 2, HPV16/59 DNA-positive/mRNA not tested; Patient 3, HPV16 DNA-positive/mRNA-negative; Patient 4, HPV52 DNA-positive/mRNA-positive; Patient 5, HPV59 DNA-positive/mRNA-positive; Patient 6, HPV56/59 DNA-positive/mRNA-positive; and Patient 7, HPV58 DNA-positive/mRNA-positive. The numbers of slides prepared from the right/left tonsils were 1,215/1,275 for Patient 1, 886/768 for Patient 2, 970/948 for Patient 3, 934/1,145 for Patient 4, 1,345/1,223 for Patient 5, 1,195/1,352 for Patient 6, and 1,233/1,045 for Patient 7. All seven patients were male and underwent tonsillectomy for recurrent tonsillitis. Patient 2 did not provide a second gargle specimen; thus, his oral HR-HPV mRNA status was unknown. Slides were assessed for HR-HPV DNA matching the gargle genotype (Extended Data Fig. 1). The number of HPV16 DNA-positive slides for the right and left tonsils was as follows: 0 of 25 and 14 of 70 for Patient 1; 11 of 18 and 0 of 22 for Patient 2; and 0 of 21 and 1 of 28 for Patient 3. No HPV59-positive slide was found in Patient 2. HPV52 DNA was detected in 0 of 19 slides and 3 of 32 slides for the right and left tonsils, respectively, in Patient 4; HPV59 DNA in 2 of 28 and 24 of 26 in Patient 5; HPV56 DNA in 3 of 27 and 0 of 29 and HPV59 DNA in 0 of 27 and 3 of 29 in Patient 6; and HPV58 DNA in 0 of 27 and 11 of 23 in Patient 7. Viral load was high in Patients 2, 5, and 6, with extensive consecutive positivity in the left tonsil of Patient 5. For subsequent analyses, peak viral-load slides (one per peak) were selected; for clustered positive slides, the highest-load slide was selected. Each selected slide plus adjacent slides was defined as a “Site” (Extended Data Fig. 1).

### HPV16-associated intraepithelial lesions in the tonsil

HPV16-associated intraepithelial lesions with moderate-to-severe atypia were identified in Patients 1 and 2. Patient 1 had one lesion at Site 1-1 (Lesion 1-1), two at Site 1-2 (Lesions 1-2-1 and 1-2-2), and one at Site 1-3 (Lesion 1-3); Patient 2 had one lesion at Site 2-1 (Lesion 2-1). Histological and immunohistochemical findings for Lesions 1-2-1, 1-2-2, 1-3, and 2-1 are shown in Fig. 3, and those for Lesion 1-1 in Extended Data Fig. 2. The longest diameters of Lesions 1-1, 1-2-1, 1-2-2, 1-3, and 2-1 were 300, 1,400, 400, 630, and 2,500 μm, respectively.

**Fig. 3:**
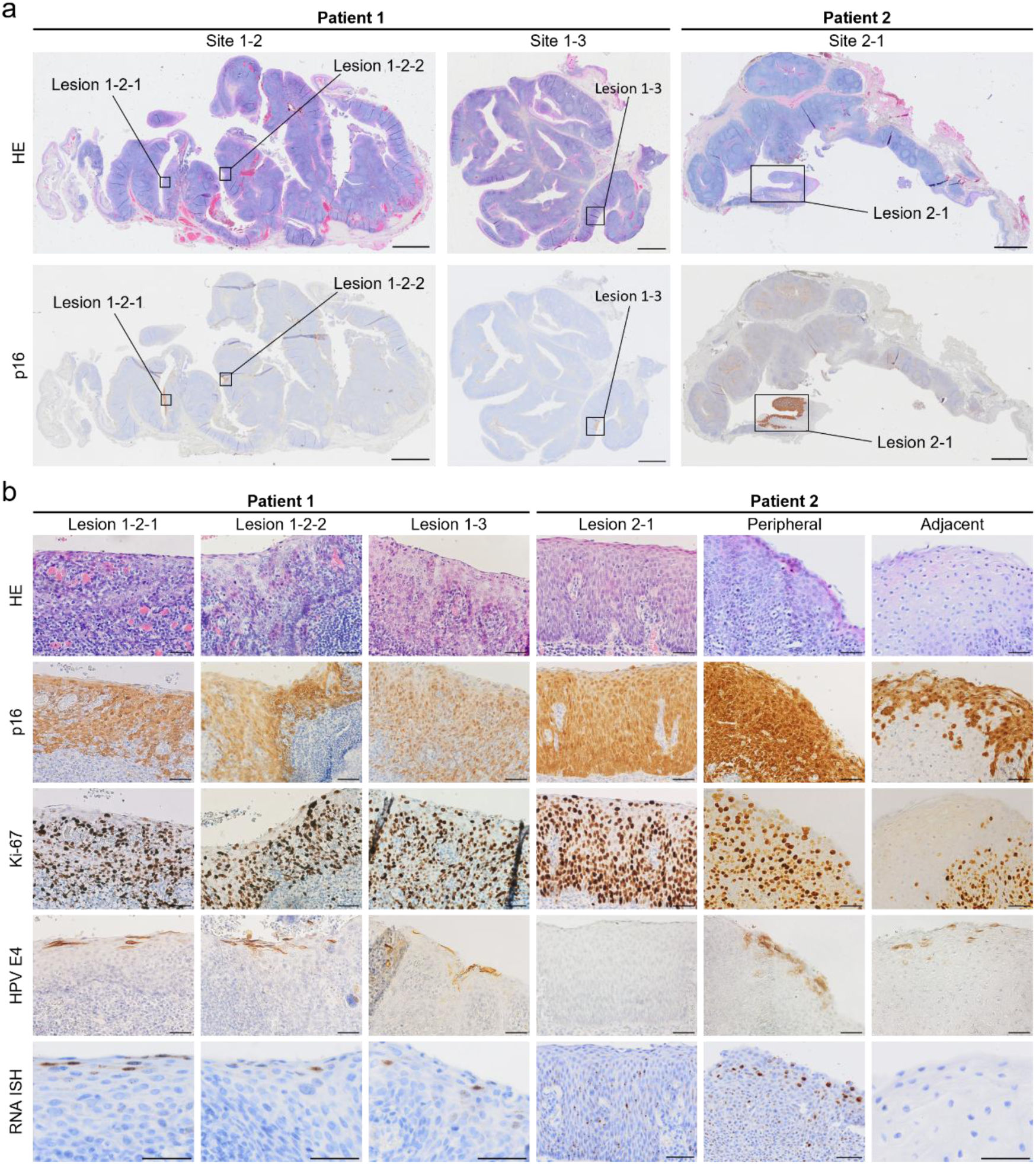
Histopathological, immunohistochemical, and *in situ* hybridization findings of HPV16-associated tonsillar intraepithelial lesions. **a.** Whole-section images of Sites 1-2 and 1-3 in Patient 1 and Site 2-1 in Patient 2. The top and bottom rows show hematoxylin and eosin (HE) staining and p16 immunohistochemistry (IHC), respectively; HPV-associated lesions are boxed. **b.** High-magnification images showing, from top to bottom, HE staining, p16 IHC, Ki-67 IHC, HPV E4 IHC, and HR-HPV E6/E7 mRNA *in situ* hybridization. Scale bars, 2 mm (a) and 50 μm (b). HPV, human papillomavirus; HR-HPV, high-risk HPV.

The tonsil parenchyma consists of non-keratinized stratified squamous epithelium and lymphoid nodules, with numerous crypts extending from the tonsillar surface deep into the parenchyma. Normal tonsillar epithelium comprises one basal cell layer, two to three parabasal layers of proliferating basaloid cells, and intermediate and superficial layers in which cells progressively flatten and the nucleus-to-cytoplasm (N/C) ratio decreases toward the surface. In contrast, HPV16-associated intraepithelial lesions showed expansion of atypical basaloid cells with high N/C ratios, enlarged nuclei, and irregular nuclear contours beyond the lower one-third of the epithelium. Markedly enlarged nuclei and mitotic figures were common, particularly in Lesions 1-2-2 and 2-1. Surface maturation was preserved, with progressive squamous flattening toward the lumen, distinguishing these lesions from full-thickness atypia.

All lesions showed block positivity for p16 and Ki-67 throughout the epithelium, whereas normal epithelium was negative or focally positive for p16 and expressed Ki-67 only in the basal and parabasal layers. HPV E4 and HR-HPV mRNA patterns varied among lesions. In Lesions 1-1, 1-2-1, 1-2-2, and 1-3, HPV E4 was confined to the superficial layer; punctate nuclear and cytoplasmic HR-HPV mRNA signals were scattered throughout the epithelium, with clustered nuclear signals in superficial layers. Lesion 2-1 lacked HPV E4 expression and showed densely distributed punctate HR-HPV mRNA signals without superficial clustering. Because Lesion 2-1 was relatively large, we also examined its peripheral and adjacent areas. The peripheral area showed atypia similar to the central area, but an expression profile resembling Lesions 1-1, 1-2-1, 1-2-2, and 1-3: block positivity for p16 and Ki-67, superficial HPV E4 expression, and clustered superficial HR-HPV mRNA signals. In the adjacent area, p16 expression was focal and Ki-67 was confined to basal and parabasal layers. No HR-HPV mRNA signal was detected, but HPV E4-positive cells with a koilocyte-like appearance were present. Histological changes across Lesion 2-1 coincided with changes in p16, Ki-67, HPV E4, and HR-HPV mRNA expression patterns.

At Site 2-2 in Patient 2, no atypical lesion or p16 or HPV E4 expression was observed; however, a lesion exhibiting sporadic HR-HPV mRNA expression was identified (Lesion 2-2; Extended Data Fig. 2). Lesion 2-2 expressed Ki-67 throughout the epithelium. Site 2-3 in Patient 2 and Site 3-1 in Patient 3 showed normal histology and Ki-67 expression and no p16 or HPV E4 expression (data not shown).

### Intraepithelial lesions in the tonsil associated with high-risk HPV other than HPV16

Multiple HPV59-associated intraepithelial lesions with mild atypia were identified at Sites 5-1 and 5-2 in Patient 5. In Patient 6, a solitary HPV56-associated intraepithelial lesion with minimal atypia was identified at Site 6-1 of the right tonsil (Lesion 6-1), and a similar HPV59-associated lesion at Site 6-2 of the left tonsil (Lesion 6-2). Representative lesions at Site 5-2 (Lesions 5-2-1 and 5-2-2) and Lesions 6-1 and 6-2 are shown in Fig. 4. Atypical cells were confined to the lower third of the epithelium. Ki-67 was expressed throughout the epithelium, whereas p16 was predominantly expressed in the lower two-thirds. HPV E4 was detected in the upper third, and punctate signals of HR-HPV mRNA were widely observed in the basal and parabasal layers, with strongly clustered nuclear signals in the upper epithelium. Lesions 5-2-1, 5-2-2, 6-1, and 6-2 measured 2,600, 480, 570, and 450 μm in the longest diameter, respectively.

**Fig. 4:**
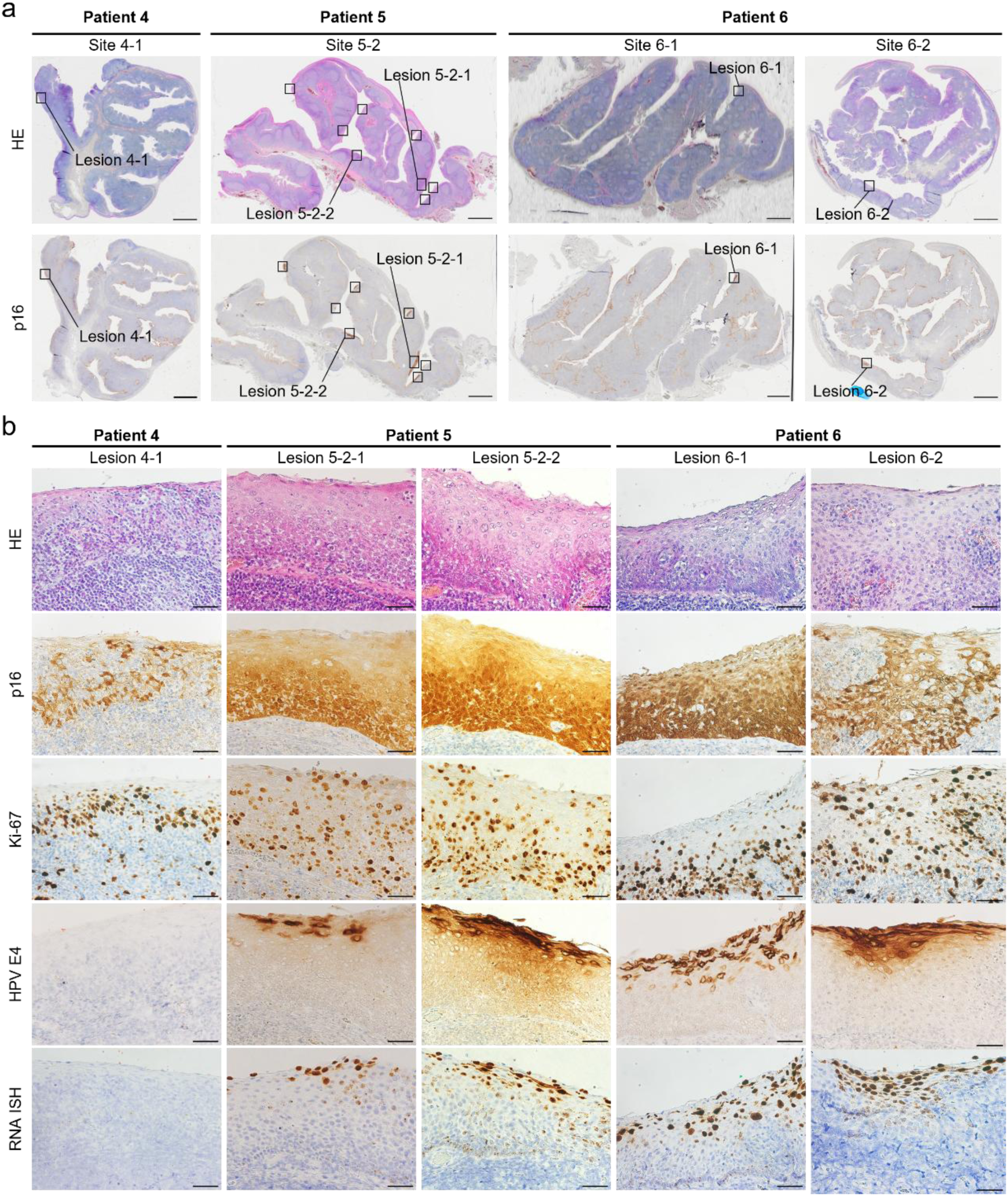
Histopathological, immunohistochemical, and *in situ* hybridization findings of tonsillar intraepithelial lesions associated with high-risk HPV other than HPV16. **a.** Whole-section images of Site 4-1 in Patient 4, Site 5-2 in Patient 5, and Sites 6-1 and 6-2 in Patient 6. The top and bottom rows show hematoxylin and eosin (HE) staining and p16 immunohistochemistry (IHC), respectively; HPV-associated lesions are boxed. **b.** High-magnification images showing, from top to bottom, HE staining, p16 IHC, Ki-67 IHC, HPV E4 IHC, and HR-HPV E6/E7 mRNA *in situ* hybridization. Representative lesions are shown for Site 5-2, which contained multiple HPV-associated lesions. Scale bars, 2 mm (a) and 50 μm (b). HPV, human papillomavirus; HR-HPV, high-risk HPV.

No atypical lesions were identified at Site 4-1 in Patient 4. However, Lesion 4-1 showed patchy p16 positivity throughout the epithelium and Ki-67 negativity in superficial layers (Fig. 4). HR-HPV mRNA signals were restricted to a 650-μm-wide region of the basal layer (Extended Data Fig. 3), and HPV E4 was absent throughout the epithelium. At Site 7-1 in Patient 7, histology and p16 and Ki-67 expression were comparable to normal epithelium, and neither HPV E4 nor HR-HPV mRNA was detected (data not shown). Slides adjacent to all other HPV58 DNA-positive slides showed similar findings (data not shown).

Table 1 summarizes HPV test results and HPV-associated lesions in the seven patients whose tonsils were examined. Intraepithelial lesions with atypia were identified in three of five patients positive for oral HR-HPV mRNA and in one patient with oral HPV16 infection whose oral HR-HPV mRNA status was not tested. All lesions were confined to tonsillar crypts.

**Table 1.**
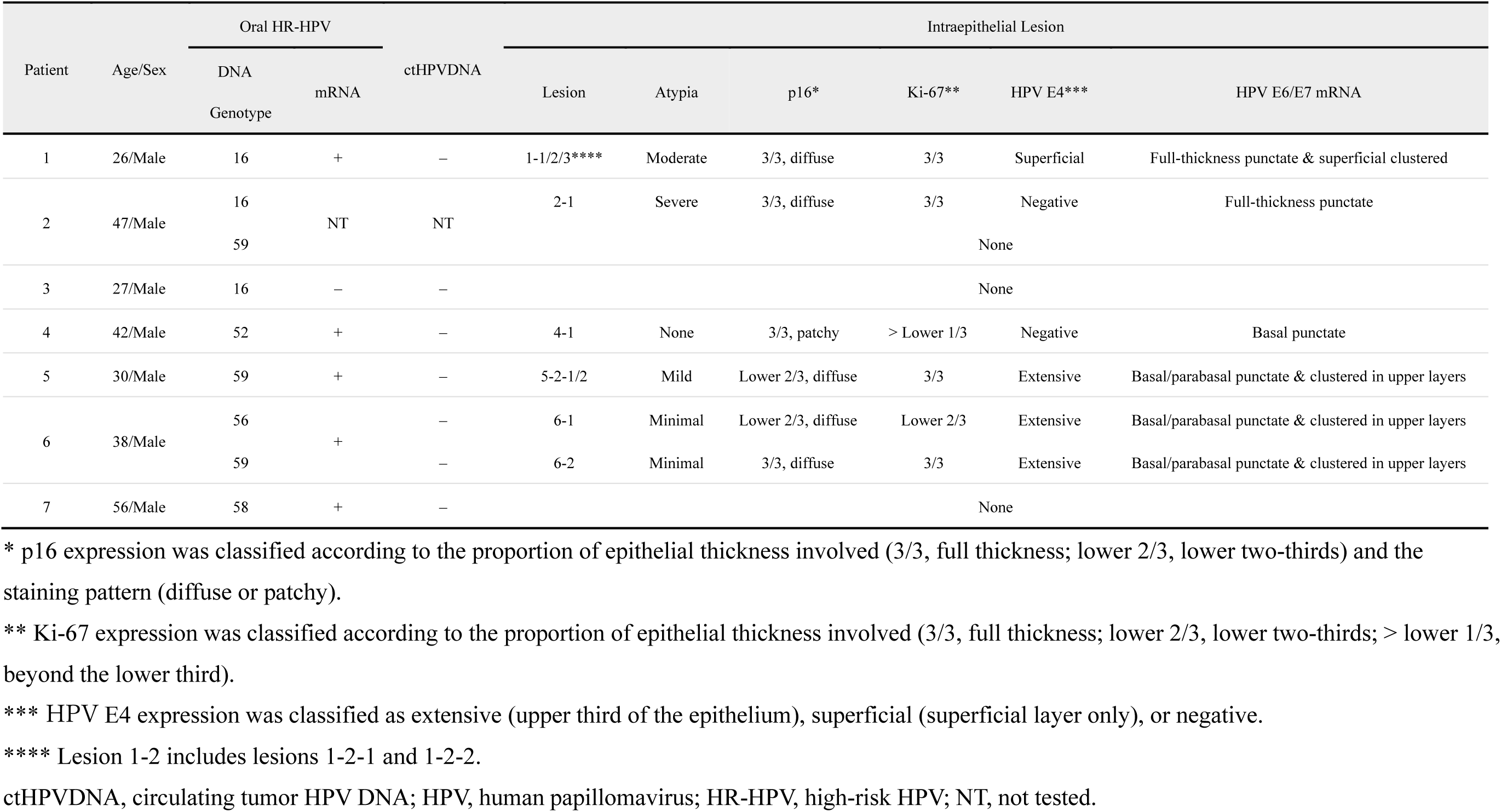
Summary of HPV tests and findings on HPV-associated intraepithelial lesions.

### Spatial molecular profiles of HPV-associated intraepithelial lesions

Tissue sections containing Lesions 2-1, 5-2, and 6-2 met the quality threshold (DV200 > 30%) and proceeded to Visium HD analysis, whereas the sections containing Lesions 1-1, 1-2, 1-3, 2-2, 4-1, and 6-1 had suboptimal RNA quality (DV200 < 30%) and were not analyzed. Based on morphological severity, Lesions 5-2-1 and 6-2 were classified as low-grade dysplasia, and Lesion 2-1 as high-grade dysplasia.

Using public single-cell datasets, we deconvolved 8-µm Visium HD bins to spatially annotate predominant cell types (Fig. 5a). For molecular-transition analysis, we retained high-purity epithelial bins (>80% epithelial content); numbers and proportions of retained bins are shown in Extended Data Table 6. An integrated uniform manifold approximation and projection (UMAP) of these bins, combined with expert pathological annotations and trajectory inference, revealed a continuous molecular shift from normal epithelium through low- and high-grade dysplasia to invasive cancer (Fig. 5b). Pseudotime distributions across pathological states further supported this continuum (Extended Data Fig. 4).

**Fig. 5:**
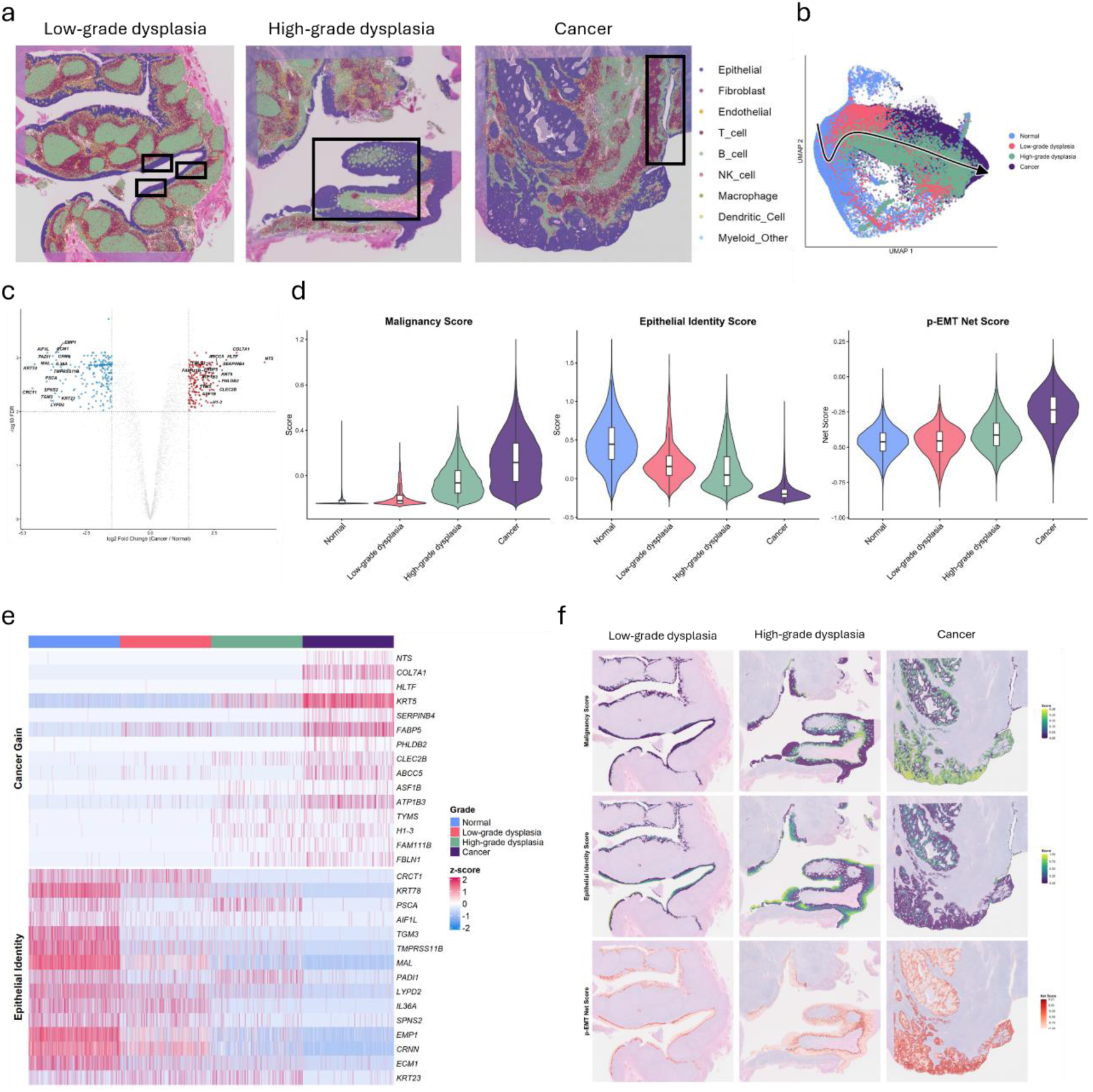
Spatial transcriptomic profiles support stepwise HPV-associated carcinogenesis in the tonsil. **a.** Spatial cell-type deconvolution of representative Visium HD sections across histological grades. Bins are color-coded by predominant cell type. Low-grade dysplasia, high-grade dysplasia, and cancer are represented by Lesion 5-2-1 (Patient 5), Lesion 2-1 (Patient 2), and OPC1, respectively. Dysplastic lesions are boxed; two additional lesions are present near Lesion 5-2-1. In the cancer section, normal epithelium is boxed because cancer occupies most of the section. OPC1 is described in Extended Data Table 5. **b.** UMAP projection of epithelial-rich bins (purity >80%) integrated across six sections, showing a molecular continuum from normal epithelium through dysplasia to cancer. The black arrow indicates the inferred progression trajectory. **c.** Volcano plot of differentially expressed genes (DEGs) between normal epithelium (n=6) and cancer (n=3). Genes with FDR <0.01 and |log2FC| >1.5 are shown in red (upregulated) or blue (downregulated). **d.** Malignancy, epithelial identity, and p-EMT Net scores across histological grades (normal, n=6; low-grade dysplasia, n=2; high-grade dysplasia, n=1; cancer, n=3). Boxes show the interquartile range (IQR), center lines the medians, and whiskers 1.5 × IQR. **e.** Heatmap of the top 15 DEGs ranked by log2FC, showing Z-score-normalized expression across normal epithelium, low-grade dysplasia, high-grade dysplasia, and cancer. **f.** Spatial mapping of the scores in **d** for the lesions shown in **a**. DEG, differentially expressed gene; FDR, false discovery rate; IQR, interquartile range; log2FC, log2 fold change; p-EMT, partial epithelial-mesenchymal transition; UMAP, uniform manifold approximation and projection.

Differentially expressed genes (DEGs) between normal and malignant epithelia defined signatures of normal differentiation and malignancy (Fig. 5c), from which we calculated malignancy, epithelial identity, and partial epithelial-mesenchymal transition (p-EMT) Net scores. All three scores varied with histological grade from normal epithelium through low- and high-grade dysplasia to cancer (Fig. 5d). A heatmap of the top 30 DEGs further showed the intermediate features of dysplastic lesions (Fig. 5e): low-grade dysplasia resembled normal tissue for both “Cancer-gain” and “Epithelial-identity” genes, whereas high-grade dysplasia upregulated a subset of “Cancer-gain” genes while retaining near-normal “Epithelial-identity” expression. Spatial plots were concordant with the histological and molecular changes (Fig. 5f). Malignancy and p-EMT Net scores increased throughout the epithelial layers in cancer but only in basal and parabasal layers in high-grade dysplasia. The epithelial identity score remained near normal in the upper third of high-grade dysplasia, whereas low-grade dysplasia more closely resembled normal epithelium. Overall, HPV-associated intraepithelial lesions with atypia had molecular profiles intermediate between normal and malignant epithelium.

## DISCUSSION

In this study, we identified HPV-associated tonsillar intraepithelial lesions that were histologically dysplastic and biologically distinct from both normal epithelium and HPV-associated oropharyngeal cancer. These lesions were likely precursors of HPV-associated oropharyngeal cancer, and we found two distinct patterns. One was intraepithelial lesions with moderate-to-severe atypia, showing block positivity for p16 and Ki-67 throughout the epithelium; HPV E4 was negative or restricted to the superficial layer, and HR-HPV E6/E7 mRNA was distributed throughout the epithelium with or without superficial clustering. The other was intraepithelial lesions with minimal-to-mild atypia, showing p16 and Ki-67 predominantly in the lower two-thirds of the epithelium, high expression of HPV E4 in the upper third, and HR-HPV E6/E7 mRNA distributed in the lower two-thirds with strong clustering in the upper third. Tonsillar intraepithelial lesions might therefore be graded by dysplasia severity, analogous to CIN. However, tonsillar crypt epithelium differs architecturally and immunologically from cervical squamous epithelium^20^; thus, CIN grading could not be directly applied. A tonsil-specific grading system will require analysis of substantially more cases.

In terms of HPV life cycle, CIN is categorized into productive, intermediate, and transforming lesions according to HPV E4, p16, and Ki-67 expression patterns.^10,34–36^ Following this categorization, tonsillar lesions with moderate-to-severe atypia (Lesions 1-1, 1-2-1, 1-2-2, 1-3, and 2-1) were transforming lesions, whereas those with minimal-to-mild atypia (Lesions 5-2-1, 5-2-2, 6-1, and 6-2) were productive-to-intermediate lesions. The absence of HPV E4 in Lesion 2-1 suggests a transforming lesion close to progression to cancer. Within Lesion 2-1, the central area was transforming, the peripheral area retained productive activity despite a transforming phenotype, and the adjacent area was productive, forming a continuous sequence of HPV life-cycle stages. Spatial transcriptomics likewise showed a molecular continuum from normal epithelium through atypical intraepithelial lesions to invasive cancer. Näsman et al. similarly reported sequential progression from adjacent dysplasia to invasive HPV-associated oropharyngeal cancer by trajectory analysis.^37^ Together, these findings support stepwise HPV-driven tonsillar carcinogenesis, although its natural history remains incompletely defined. Lesion 4-1, characterized by weak basal HR-HPV mRNA expression and absent HPV E4, might represent latent HPV infection.^38^ Reactivation of latent HPV could therefore contribute to some HPV-associated oropharyngeal cancers, although latency and reactivation have not been reported in the tonsils and are supported only indirectly even in the cervix.^10,39^ HPV16-associated lesions showed moderate-to-severe atypia, whereas lesions associated with other HR-HPV types showed minimal-to-mild atypia. Although based on very few cases, this difference is notable because HPV16 causes most HPV-associated oropharyngeal cancers. HPV16-associated lesions may be more likely to evade immune clearance and progress, whereas lesions associated with other HR-HPV types may be more likely to regress. The higher frequency of oral HR-HPV E6/E7 mRNA among patients with persistent HPV16 infection than among those with persistent infection by other HR-HPV types may further support this possibility.

As expected, all HPV-associated intraepithelial lesions were confined to the tonsillar crypts rather than to the tonsillar surface. Therefore, direct visualization of HPV-associated early tonsillar lesions is impractical, underscoring the need for biomarkers for early detection. Candidates include HPV DNA in oral rinses or gargles, HPV16 E6 antibody in plasma or serum, and ctHPVDNA.^11^ Each biomarker has limitations. Oral HPV DNA indicates infection but does not necessarily indicate the presence of HPV-associated lesions. Most oral HPV DNA clears within one year,^25^ and subsequent cancer risk after oral HPV DNA positivity is very low.^40^ In contrast, the HPV16 E6 antibody is stable. Less than 1% of healthy individuals are seropositive for HPV16 E6 antibody, but most seropositive individuals do not develop HPV-associated oropharyngeal cancer; limiting its utility alone.^41,42^ Because ctHPVDNA levels correlate with tumor burden and HPV copy number per tumor cell,^30^ ctHPVDNA is unlikely to be a surrogate for HPV-associated early lesions. In fact, ctHPVDNA was not detected in any patient with HPV-associated intraepithelial lesions in this study. Therefore, other biomarkers are needed for the early detection of HPV-associated oropharyngeal cancer.

Oral HR-HPV mRNA was detected in three of four patients with HPV-associated intraepithelial lesions with atypia. The remaining patient did not undergo oral HR-HPV mRNA testing, but his lesion contained abundant HR-HPV mRNA, suggesting that oral testing would likely also have been positive. These findings support oral HR-HPV mRNA as a candidate biomarker for HPV-associated early lesions. However, oral HR-HPV mRNA testing was restricted to patients with oral HR-HPV infection. In this study, no patient without persistent oral HR-HPV infection had detectable oral HR-HPV mRNA, and a separate study likewise detected no oral HR-HPV mRNA in healthy individuals without oral HR-HPV infection.^31^ Thus, oral HR-HPV mRNA is unlikely to be detectable without oral HR-HPV infection. Because serial sectioning to identify latent tonsillar lesions is extremely labor-intensive, tonsillar examination was limited to patients positive for oral HR-HPV mRNA and those with oral HPV16 infection regardless of mRNA status. We therefore could not estimate the sensitivity or specificity of oral HR-HPV mRNA for HPV-associated early lesions, but the present findings provide a rationale for further studies. We have introduced an automated serial sectioning system and initiated a cohort study of patients undergoing tonsillectomy for benign conditions in which preoperative gargles from all participants will be tested for HR-HPV DNA and mRNA. This cohort should enable the evaluation of the sensitivity and specificity of oral HR-HPV mRNA as a biomarker for HPV-associated early lesions.

Among the five patients positive for oral HR-HPV mRNA, two had no detectable HPV-associated tonsillar lesion. Tonsillar HR-HPV mRNA was absent in one patient, consistent with infection foci without an active lesion, and was faintly detected only in the basal layer in the other, consistent with possible viral latency. Because the base of the tongue is the second most common site of HPV-associated oropharyngeal cancer after the tonsils,^43^ occult lesions at the tongue base cannot be excluded in these patients despite negative endoscopy; the same possibility applies to patients with confirmed tonsillar lesions. A major limitation of tonsillectomy-based studies is that the tonsils can be examined exhaustively whereas the tongue base cannot. In our ongoing cohort, oral gargles will therefore be tested for HR-HPV mRNA before and after tonsillectomy. Persistent positivity after tonsillectomy could indicate an occult lesion at the tongue base. Such patients will undergo long-term follow-up with periodic oral HR-HPV mRNA assessment, during which a tongue-base lesion may become clinically apparent.

There are some other limitations in this study. The Aptima HPV assay for HR-HPV mRNA detection has been fully validated for cervical cancer screening but is not optimized for oral rinses or gargles, and the collection procedure for oral specimens is not standardized. These issues do not affect the present findings but must be resolved before oral HR-HPV mRNA can be established for early screening of HPV-associated oropharyngeal cancer.

In conclusion, we provide the first evidence of HPV-associated premalignant lesions in the tonsils and identify oral HR-HPV mRNA as a candidate biomarker. The regression, persistence, and progression rates of these lesions and other aspects of the natural history of oropharyngeal HPV infection remain unknown. Further studies are needed to define this natural history, establish oral HR-HPV mRNA as a biomarker of HPV-associated oropharyngeal early lesions, and ultimately enable early detection of HPV-associated oropharyngeal cancer. Given their analogy to CIN, we propose the term tonsillar intraepithelial neoplasia (TIN) for these lesions.

## METHODS

### Workflow for identifying HPV-associated lesions in the tonsils

This study included patients undergoing tonsillectomy for benign conditions at the University of Osaka Hospital and affiliated hospitals (Extended Data Table 7). Oral gargle specimens were collected in an outpatient setting for HPV DNA detection and genotyping (first test). Immediately before surgery, a second gargle specimen was collected for oral HPV DNA detection and genotyping and HR-HPV mRNA testing (second test), but only from patients positive for oral HR-HPV DNA on the first test. Blood was also collected from these patients for ctHPVDNA testing. Resected tonsils were formalin-fixed, paraffin-embedded (FFPE), serially sectioned in full, and mounted on slides.

DNA was extracted from every 50 slides, and droplet digital PCR (ddPCR) was used to identify slides containing HR-HPV DNA matching the genotype detected in gargle specimens. Adjacent slides were examined by hematoxylin and eosin (HE) staining, immunostaining for p16, Ki-67, and HPV E4, and *in situ* hybridization for HR-HPV mRNA to identify HPV-associated lesions. HPV-associated intraepithelial and malignant lesions were additionally analyzed by single-cell-scale spatial transcriptomics.

The study was approved by the Institutional Review Board of each participating hospital and registered with UMIN-CTR (UMIN000036583). All patients provided written informed consent. Because the age of adulthood in Japan was 20 years until April 1, 2022, patients aged <20 years, for whom parental consent would have been required, were excluded.

### Assessment of oral HR-HPV DNA and mRNA

Patients were requested to gargle with 10 mL of saline for 30 s, and the specimen was collected in a sterile collection tube for both tests. Two milliliters of each gargle specimen from the second test were aliquoted into Aptima urine specimen transport tubes (Hologic, Inc., San Diego, CA, USA). Both specimen types were immediately shipped at 4 °C to LSI Medience Corporation (Tokyo, Japan) and stored at 4 °C until analysis. DNA was extracted from collection-tube specimens using QIAsymphony SP/AS Instruments (Qiagen, Hilden, Germany) and analyzed with the GENOSEARCH HPV31 kit (Medical and Biological Laboratories, Nagoya, Japan) for HPV detection and genotyping, as previously described.^30,44^ The GENOSEARCH HPV31 kit detects 31 genotypes: high-risk types 16, 18, 31, 33, 35, 39, 45, 51, 52, 56, 58, 59, and 68; intermediate-risk types 26, 53, and 66; and low-risk types 6, 11, 42, 44, 54, 55, 61, 62, 70, 71, 73, 82, 84, 90, and CP6108. HPV mRNA was assessed using the Aptima HPV assay and Panther system (Hologic). The Aptima assay detects E6/E7 mRNA from 14 HR-HPV genotypes (HPV16, 18, 31, 33, 35, 39, 45, 51, 52, 56, 58, 59, 66, and 68) without genotype-specific identification. Although HPV66 is classified as high risk by the Aptima assay, it is generally categorized as intermediate risk and was not classified as high risk in this study.

### Measurement of circulating tumor HPV DNA

Blood samples (8.5 mL) were collected in cell-free DNA collection tubes (Roche, Basel, Switzerland) and shipped to the University of Osaka. Following centrifugation at 3,500 rpm for 10 min, plasma was collected and stored at −80 °C until use. Cell-free DNA was extracted using the QIAamp Circulating Nucleic Acid Kit (Qiagen), and ctHPVDNA matching the genotype detected in the oral gargle specimen was measured. For HPV16, E6 and E7 copy numbers were quantified by ddPCR using the QX200 Droplet Digital PCR System (Bio-Rad, Hercules, CA, USA), as previously reported.^30,45,46^ For non-HPV16 HR-HPV, either E6 or E7 copy number was quantified by ddPCR as previously described.^47–49^ Each reaction contained 5 µL extracted cell-free DNA per well, and all samples were analyzed in duplicate. The human RPP30 gene (dHsaCP2500350; Bio-Rad) served as an endogenous control for extraction efficiency and DNA quality. Target HPV probes were labeled with 6-carboxyfluorescein and the RPP30 probe with hexachlorofluorescein. Each run included appropriate positive and negative controls. Droplets were read using the QX200 Droplet Reader (Bio-Rad), and data were analyzed using the QuantaSoft v1.7.4.0917 (Bio-Rad). Primer and probe sets are listed in Extended Data Table 8.

### Detection of HPV DNA-positive sites in the tonsil

Excised tonsils fixed in 10% formalin were shipped to the University of Osaka, divided into two or three pieces, and paraffin embedded. Serial 6-µm sections were prepared from the entire tonsils using a manual microtome (Leica Biosystems, Nussloch, Germany). Depending on tissue size, one to four consecutive sections were placed on each slide. Slides were stretched at 42 °C, dried at 60 °C for several hours, and stored at 4 °C until use. DNA was extracted from every 50 slides using the QIAamp DNA FFPE Tissue Kit (Qiagen, Hilden, Germany). DNA concentration was measured with a Qubit 2.0 fluorometer and Qubit dsDNA BR Assay Kit (Invitrogen, Carlsbad, CA, USA), and samples >20 ng/µL were diluted to 20 ng/µL. Each ddPCR well contained 5 µL DNA template, with a maximum input of 100 ng per reaction. HPV copy numbers per slide were quantified with genotype-specific ddPCR assays matching the genotypes detected in gargle specimens, as described above; primer and probe sets are listed in Extended Data Table 8. A single positive droplet was considered background to exclude stochastic noise. Slides with ≥2 droplets were defined as HPV DNA-positive based on the 95% limit of detection from our previous Probit analysis of a similar ddPCR assay.^46^ When a slide contained ≥2 but <10 droplets, DNA was additionally extracted from every 10 surrounding slides for further ddPCR analysis as appropriate.

### Immunostaining for p16, Ki-67, and HPV E4 and in situ hybridization for HPV mRNA

Slides adjacent to HPV DNA-positive slides were subjected to p16, Ki-67, and HPV E4 immunostaining and HR-HPV mRNA *in situ* hybridization, as previously reported.^36,50,51^ For immunostaining, primary antibodies were p16 INK4A (JC8) mouse monoclonal antibody (Santa Cruz Biotechnology, Santa Cruz, CA, USA), Ki-67 (D2H10) rabbit monoclonal antibody (Cell Signaling Technology, Danvers, MA, USA), and Pan-HPV E4 mouse monoclonal antibody (provided by Dr. John Doorbar, University of Cambridge, Cambridge, UK). HR-HPV E6/E7 mRNA was detected by RNA in situ hybridization using cocktail RNAscope probes (Advanced Cell Diagnostics Inc., Hayward, CA, USA) recognizing 18 HR-HPV genotypes (HPV16, 18, 26, 31, 33, 35, 39, 45, 51, 52, 53, 56, 58, 59, 66, 68, 73, and 82) without genotype-specific identification. Signals for both immunostaining and RNA in situ hybridization were visualized with 3,3’-diaminobenzidine.

### Spatial transcriptomics using Visium HD

We analyzed 6-μm FFPE sections containing HPV-associated intraepithelial lesions and sections from three known HPV-associated oropharyngeal cancers. All three cancers showed diffuse p16 immunohistochemical positivity and HPV16 DNA positivity by PCR in pretreatment biopsy specimens, and all patients underwent upfront surgery without neoadjuvant therapy. The characteristics of these patients are shown in Extended Data Table 5. The quality of the sections was prescreened using the DV200 index. Sections were deparaffinized, HE-stained, and imaged, followed by *in situ* hybridization using the Visium HD Spatial Gene Expression kit (v1 Chemistry) and the Visium Human Transcriptome Probe Set v2.0 using the Visium CytAssist instrument (10x Genomics, Pleasanton, CA, USA). Libraries were sequenced using the DNBSEQ-G400RS platform. Raw sequencing data were processed using Space Ranger v4.0.1 (10x Genomics) for demultiplexing, alignment to the GRCh38 human reference genome, and spatial mapping. Spatial quality control retained spots with 50–6,000 feature counts and a mitochondrial read fraction of < 10%. To preserve biologically relevant regions with low transcriptional activity, including terminally differentiated or virus-associated intraepithelial lesions, the lower feature-count threshold was optimized to nFeature_Spatial > 50 (Extended Data Fig. 5).

### Single-cell RNA-seq reference and annotation

For high-resolution deconvolution, we used the public scRNA-seq dataset of HPV-positive head and neck cancers (GEO: GSE182227).^52^ We included seven HPV-positive tonsillar cancer samples (OPSCC6-CD45+, OPSCC6-CD45-, OPSCC9-CD45+, OPSCC9-CD45-, OPSCC17, OPSCC33-tumor, and OPSCC34-tumor) and adjacent normal tonsillar tissues (OPSCC33-normal and OPSCC34-normal), excluding base-of-tongue lesions to match the anatomical site of the Visium HD specimens. Doublets were removed using the hybrid scds approach (v1.22.0), which combines binary classification and co-expression logic. Cells were also excluded if they had <1,000 or >5,000 detected genes, >60,000 total UMI counts, >10% mitochondrial transcripts, <5% ribosomal transcripts, or >10% hemoglobin transcripts (Extended Data Fig. 6a). The 1,000-gene lower threshold matched the prefiltering threshold of the original dataset. Data were processed and SCTransform normalized in Seurat v5.3.0. Samples were integrated using Seurat’s canonical correlation analysis workflow to minimize batch and inter-individual effects; mixing of cells across patients and tissue types in integrated UMAP space supported successful batch reduction (Extended Data Fig. 6b). V(D)J and immunoglobulin genes were excluded from integration to avoid artificial clustering. Cell types were annotated manually using canonical markers (Extended Data Table 9) and by Azimuth mapping to the Human Cell Atlas tonsil reference (Extended Data Fig. 7a), then validated using lineage-specific marker expression (Extended Data Fig. 7b).

### Spatial deconvolution and cross-sample integration of high-purity epithelial bins

To minimize stromal and immune contamination, we performed epithelial-focused deconvolution using the Spacexr (Robust Cell Type Decomposition, [RCTD]) R package (v2.2.1) in ‘full’ doublet mode. We estimated the epithelial-cell fraction of each Visium HD bin and retained bins with >80% epithelial content. High-purity bins from all six cases were integrated using Seurat’s reciprocal principal component analysis (RPCA) workflow with SCTransform-normalized residuals to minimize batch and inter-individual effects. UMAP projections confirmed consistency of the integrated manifold across samples (Extended Data Fig. 8a, b).

### Clustering and pathological annotation

The integrated epithelial dataset was visualized using UMAP, and bins were clustered within each case to identify specific lesion areas. In consultation with an expert pathologist, the clusters were histologically annotated as Normal, Low-grade dysplasia, High-grade dysplasia, or Cancer (Extended Data Fig. 9). The case-specific annotations were mapped onto the integrated UMAP space.

### Trajectory inference analysis

To reconstruct the continuous transcriptional dynamics during malignant transformation, trajectory inference was performed on the top 20 principal components (PCA space) of the integrated epithelial bins using the Slingshot R package (v2.14.0). Histologically annotated “Normal” and “Cancer” clusters were specified as the starting root and terminal nodes, respectively, to infer a pseudotime lineage. The inferred trajectory was visualized on the UMAP embedding, and the resulting pseudotime values were compared across pathological grades.

### Pseudobulk differential gene expression analysis

Pseudobulk differential expression (DE) analysis (normal, n=6; cancer, n=3) used the edgeR quasi-likelihood pipeline (v4.4.2). Mitochondrial and ribosomal genes were excluded before analysis. To limit false discovery and pseudoreplication, counts were aggregated and analyzed with a patient-blocked generalized linear model (∼ Patient + Grade).^53–55^ Gene-wise negative-binomial dispersions were estimated with glmQLFit(robust = TRUE), and DEGs were tested with the quasi-likelihood F-test (glmQLFTest), thereby accounting for inter-individual variability and disease-background differences. For heatmap visualization, the 15 most upregulated and 15 most downregulated genes by log2FC were further filtered for log CPM >1 and spatial detection rate >3%. The heatmap used 1,000 randomly downsampled cells per grade from Z-score-normalized data.

### Weighted progression and p-EMT scoring

Malignancy and Normal Epithelial Identity scores were calculated from DEGs between normal epithelium and cancer bins (FDR <0.01, |log2FC| >1.5). Each score was the dot product of the scaled expression matrix and edgeR-derived log2FC weights. Z-score-normalized expression was used for heatmap visualization, with genes ordered by log2FC. Invasive potential was assessed using a p-EMT Net Score. The p-EMT signature was derived from the gene set defined for HPV-related oropharyngeal carcinoma.^52^ The Net Score was calculated as the p-EMT module score minus the epithelial differentiation module score using the gene sets in Extended Data Table 10.

### Statistical analysis

All computational and statistical analyses of transcriptomic data were performed in R v4.4.2. Gemini (version 3.1 Pro) was used to assist with generation of R scripts for data analysis. All AI-generated code was manually reviewed, executed, and rigorously validated by the authors against expected statistical and biological outputs. Module scores were compared, where appropriate, using the Wilcoxon rank-sum or Kruskal-Wallis test. Categorical clinical characteristics were compared using Fisher’s exact test, with odds ratios (ORs) and 95% CIs calculated as appropriate.

## Data availability

Spatial transcriptomic (Visium HD) data generated in this study will be deposited in a public database (for example, Gene Expression Omnibus [GEO]) and made publicly available upon publication. Public single-cell RNA-sequencing data were obtained from GEO accession GSE182227. Deidentified participant data generated in this study are available from the corresponding author upon reasonable request.

## Code availability

Custom code and scripts used for spatial transcriptomic analysis will be deposited in a public repository (for example, GitHub or Zenodo) and made publicly available upon publication. The custom code is provided as Supplementary Software for peer review.

## Acknowledgements

We thank Ms. Mayumi Aihara (Department of Otorhinolaryngology-Head and Neck Surgery, The University of Osaka Graduate School of Medicine), Mr. Masaharu Kohara, Ms. Takako Sawamura, Ms. Megumi Nihei, Ms. Mayumi Kawashima (Department of Pathology, The University of Osaka Graduate School of Medicine), and Ms. Yuri Terao (Center for Medical Research and Education, The University of Osaka Graduate School of Medicine) for technical assistance; Dr. Tohru Kiyono (Project for Prevention of HPV-related Cancer, Exploratory Oncology Research and Clinical Trial Center, National Cancer Center) for helpful discussions; and Dr. John Doorbar (University of Cambridge, UK) for generously providing the Pan-HPV E4 antibody.

## Osaka Study Consortium for Tonsillar Intraepithelial Neoplasia

Dr. Atsuhiko Uno (Osaka General Medical Center), Dr. Rie Ito, Dr. Miyu Saito (Osaka Rosai Hospital), Dr. Itsuki Kitayama, Dr. Ken Matsumoto, Dr. Shiro Akahani (Kansai Rosai Hospital), Dr. Sachiko Hio, Dr. Kengo Aoki, Dr. Nanao Matsukawa, Dr. Shuya Tatsuki (Yao Municipal Hospital), Dr. Hisanori Sasai, Dr. Hiroshi Nishimura, Dr. Takeshi Tsuda (National Hospital Organization Osaka National Hospital), Dr. Yasumitsu Takimoto, Dr. Kengo Nozaki, Dr. Yukinori Tsuruta, Dr. Hironori Cho (Osaka Police Hospital), Dr. Makoto Ogawa, Dr. Akihiro Nagata (Japan Community Health care Organization Osaka Hospital), Dr. Naoki Umeda, Dr. Kohei Matsuo (Suita Municipal Hospital), Dr. Yoshiyuki Ozono (Higashiosaka City Medical Center), Dr. Toshimichi Yasui, Dr. Keisuke Yamamoto (Otemae Hospital), Dr. Yoshiharu Sakata, Dr. Masaki Hayama (Hyogo Prefectural Nishinomiya Hospital), Dr. Michiko Hashimoto (Kinki Central Hospital), Dr. Takashi Shikina, Dr. Aya Nakagawa (Ikeda Municipal Hospital), Dr. Tomoya Fujisaki, Dr. Junpei Iga, Dr. Masayuki Nozawa, Dr. Hitoshi Akazawa, Dr. Miki Nagai (Sakai City Medical Center), Dr. Takashi Hirai, Dr. Ryosuke Koike (Sumitomo Hospital), Dr. Teiji Takemura (Minoh City Hospital), Dr. Takahiro Sasaki, Dr. Masashi Yamamoto, Dr. Yukiko Hanada (Osaka Habikino Medical Center) and Dr. Shinichi Miyaguchi (Kansai Medical Hospital).

## Author contributions

H.I. conceptualized the study. H.I., M.M., and H.T. developed the methodology. H.T. supervised sample collection. M.M., H.T., M.S., K.O., S.O., H.M., R.W., and T.Y. performed the investigations or analyses. K.K., D.M., and E.M. performed the pathological diagnoses. N.E. interpreted the histological dynamics of the HPV life cycle. M.M., N.S., T.K., and M.H. performed the spatial transcriptomic analysis. H.I. acquired funding. H.I. and M.M. wrote the original manuscript draft. All authors reviewed and approved the final manuscript, had full access to the study data, and had final responsibility for the decision to submit for publication.

---

**Extended Data Fig. 1:**
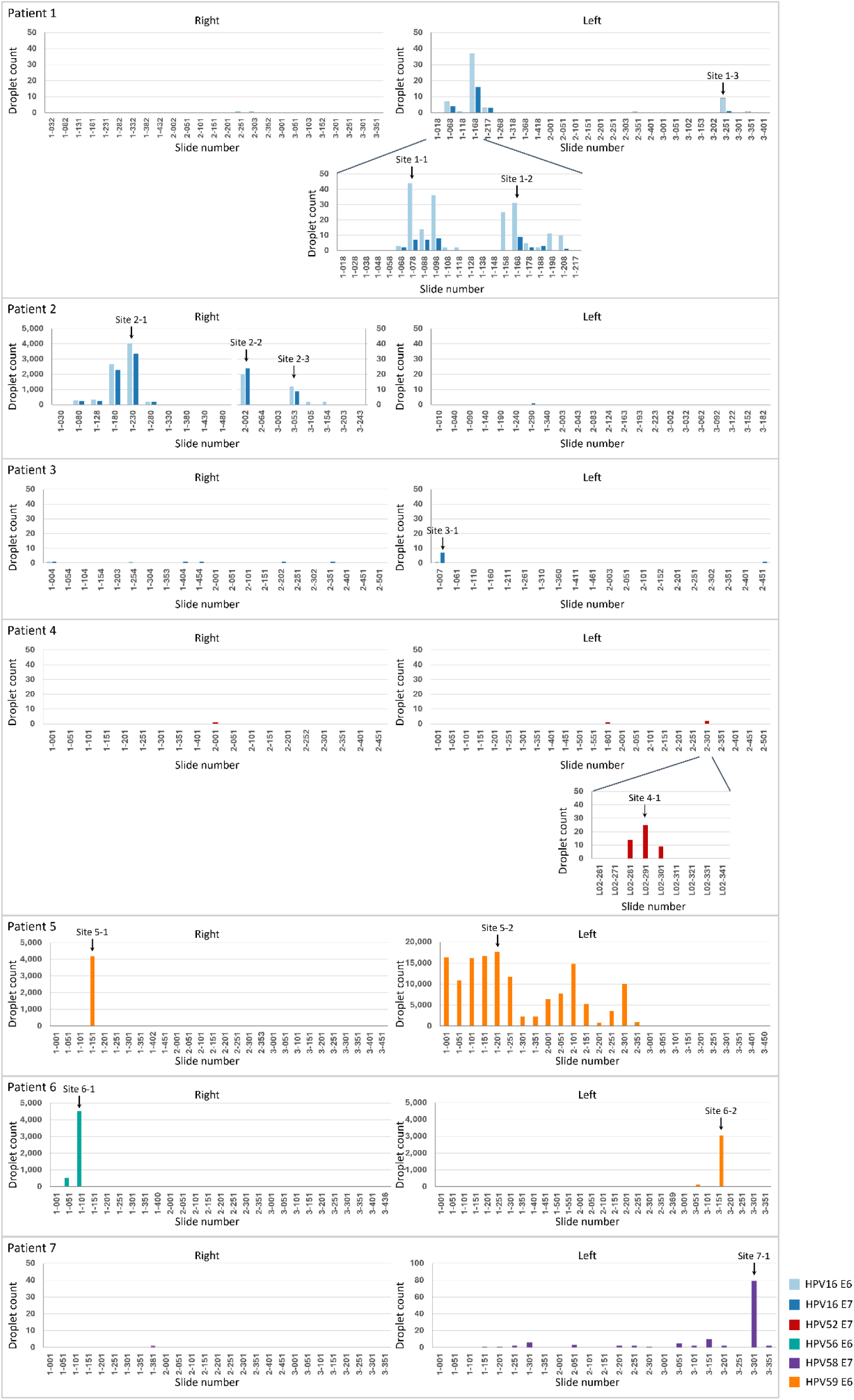
Detection of HPV DNA-positive sites in the tonsil. HPV DNA (E6 and/or E7) in each slide was quantified using ddPCR. The vertical axis shows droplet counts, with the scale varying by tonsil. For the right tonsil of Patient 2, the left and right vertical axes correspond to one-third and two-thirds, respectively, of the tonsil divided into three pieces. The horizontal axis shows slide number: the first number identifies the tonsil piece and the number after the hyphen identifies the slide within that piece. Although Patient 2 was positive for oral HPV16 and HPV59 DNA, no HPV59 DNA-positive slide was identified. Representative HR-HPV DNA-positive slides were selected for subsequent analyses, and each selected slide with its adjacent slides was designated a “Site.” ddPCR, droplet digital PCR; HPV, human papillomavirus; HR-HPV, high-risk HPV.

**Extended Data Fig. 2:**
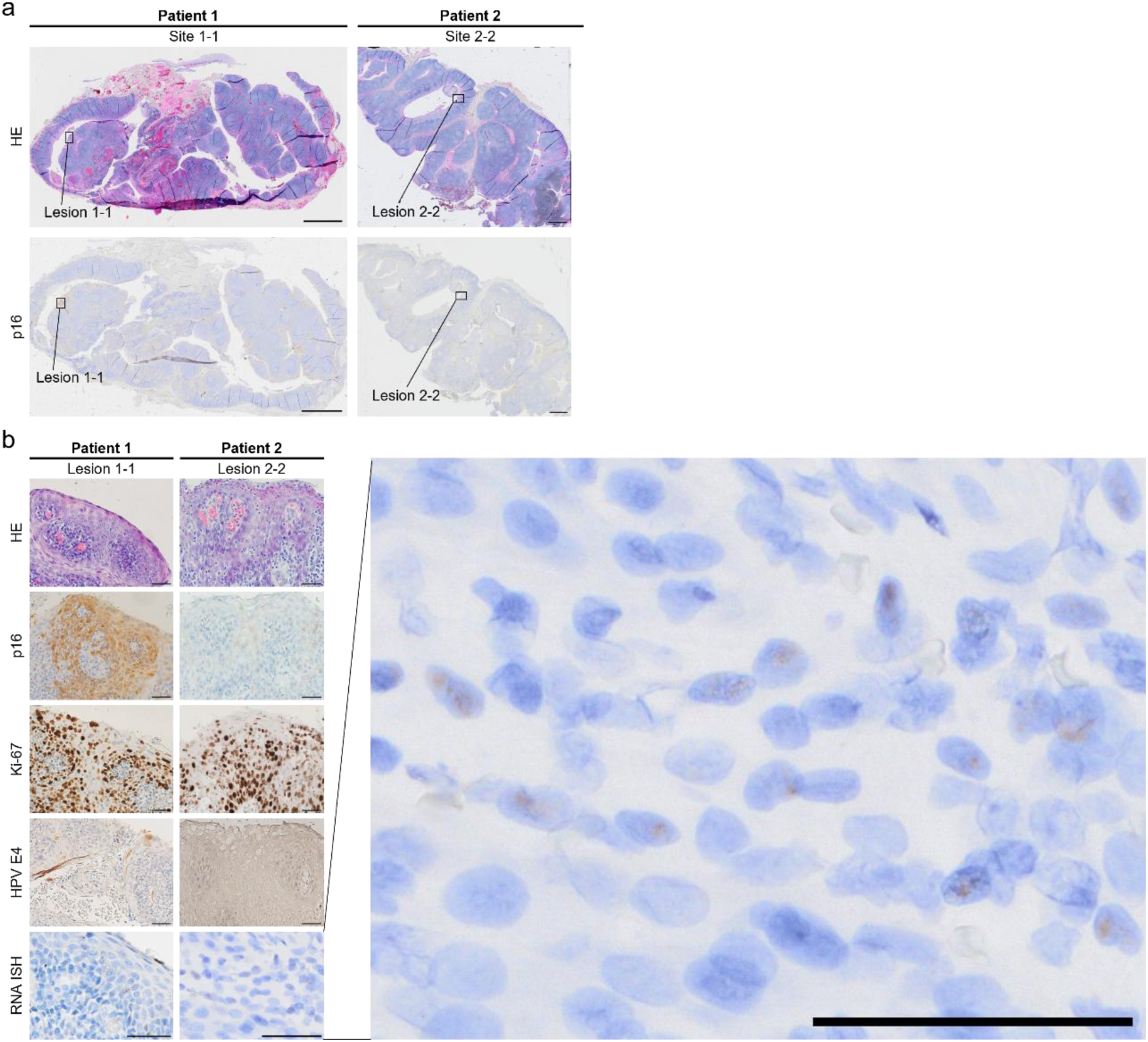
Histopathological, immunohistochemical, and *in situ* hybridization findings of HPV16-associated tonsillar intraepithelial lesions. **a.** Whole-section images of Site 1-1 in Patient 1 and Site 2-2 in Patient 2. The top and bottom rows show hematoxylin and eosin (HE) staining and p16 immunohistochemistry (IHC), respectively; HPV-associated lesions are boxed. **b.** High-magnification images showing, from top to bottom, HE staining, p16 IHC, Ki-67 IHC, HPV E4 IHC, and HR-HPV mRNA *in situ* hybridization (ISH). The rightmost panel enlarges the HR-HPV mRNA ISH image of Lesion 2-2 to show the sporadic punctate signals. Scale bars, 2 mm (a) and 50 μm (b). HPV, human papillomavirus; HR-HPV, high-risk HPV.

**Extended Data Fig. 3:**
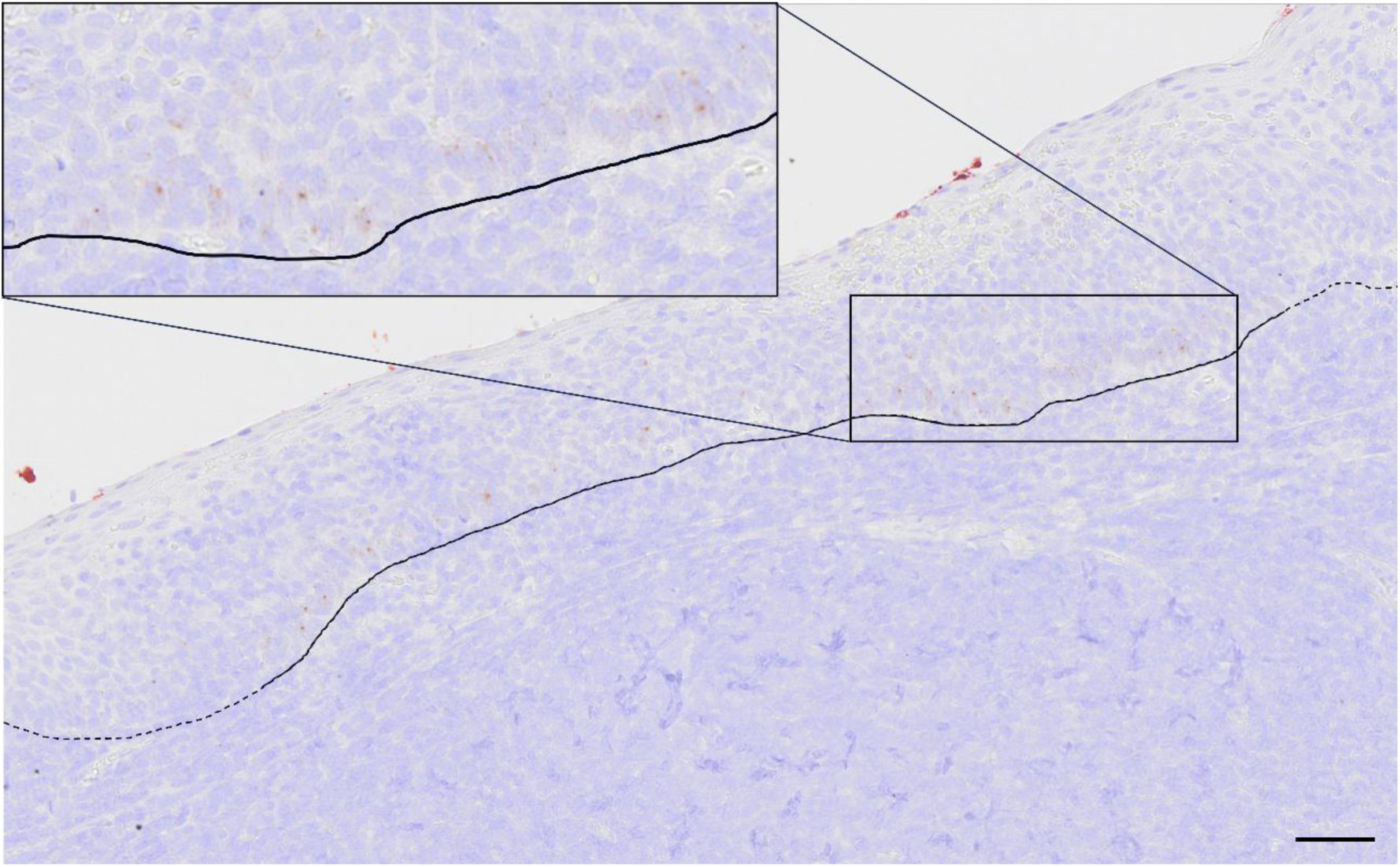
HR-HPV mRNA *in situ* hybridization in Lesion 4-1. Solid and dashed lines indicate the basement membrane. HR-HPV mRNA signals (brown punctate dots) are present in the basal layer along the solid line but absent along the dashed line. Scale bar, 50 μm. HR-HPV, high-risk human papillomavirus.

**Extended Data Fig. 4:**
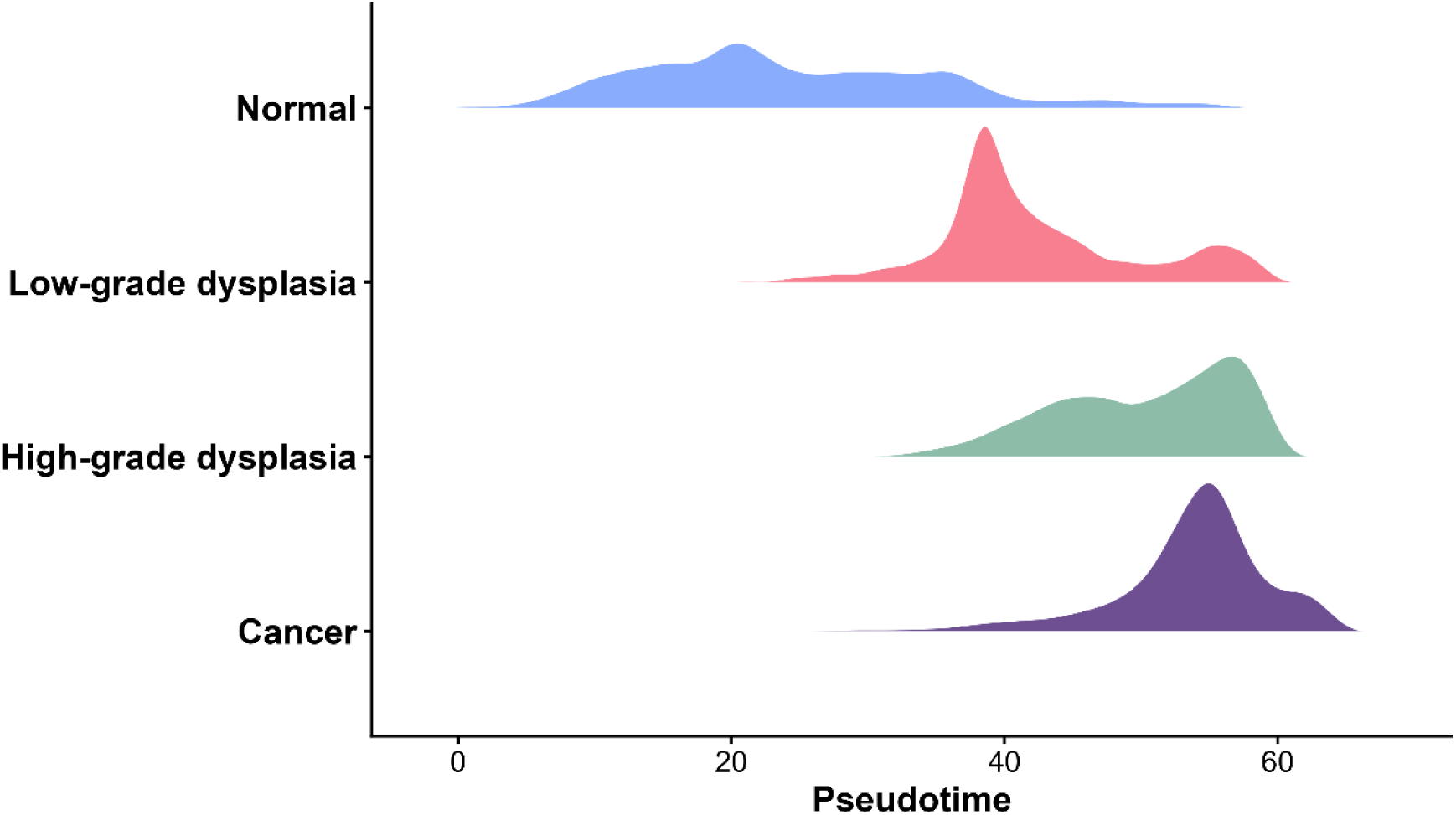
Pseudotime distribution across pathological grades. Ridge plots show the probability density distribution of pseudotime for each pathological grade. Pseudotime was calculated in PCA space. The horizontal axis shows pseudotime, and ridge height indicates spatial-bin density. The continuous stepwise shift from Normal to Cancer supports a gradual molecular transition during disease progression. PCA, principal component analysis.

**Extended Data Fig. 5:**
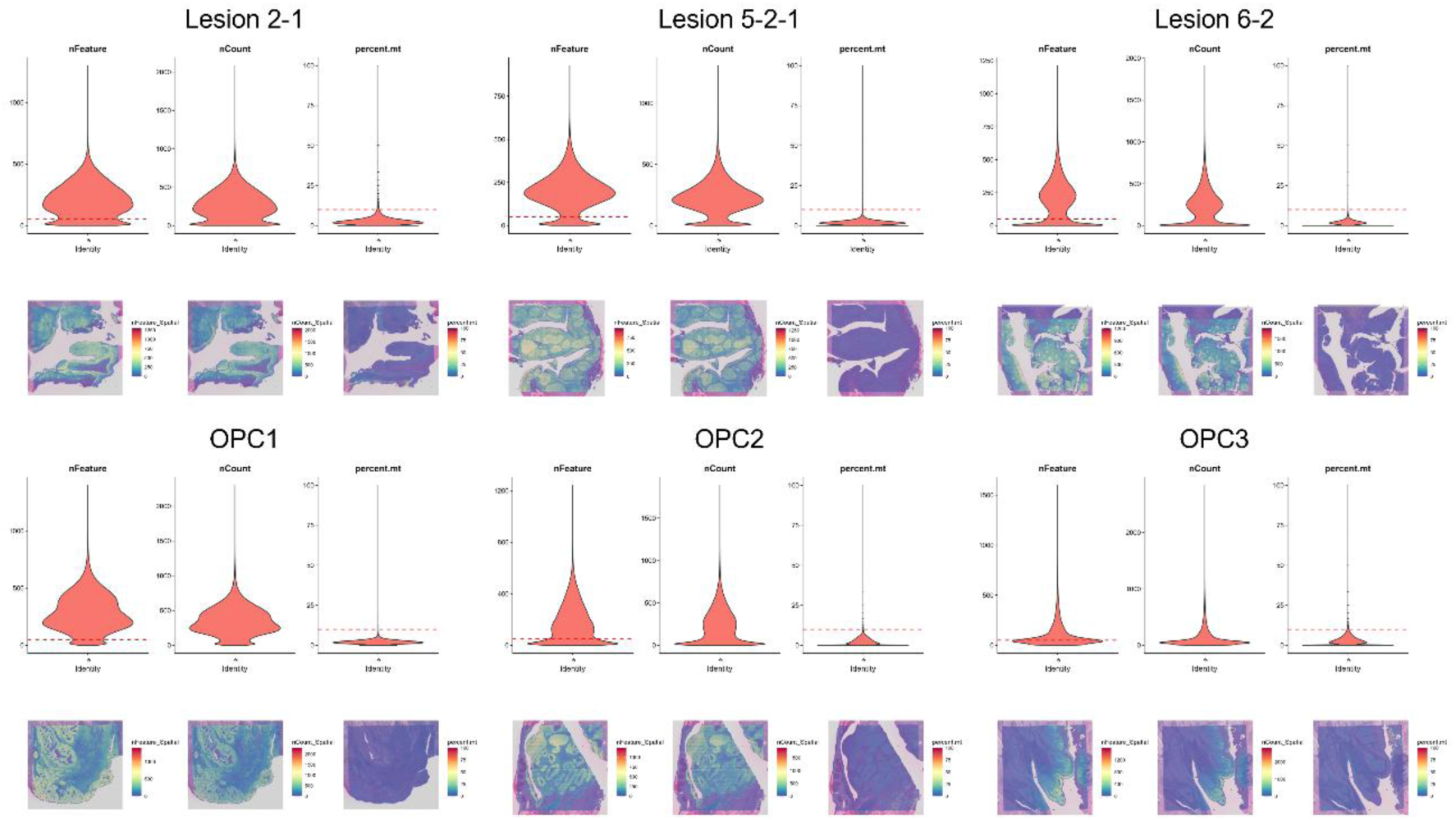
Quality control metrics and spatial distribution for Visium HD spatial transcriptomics data. For each sample (Lesions 2-1, 5-2-1, and 6-2; OPC1, OPC2, and OPC3), violin plots (top) and spatial maps (bottom) show the number of detected genes (nFeature_Spatial), total transcript counts (nCount_Spatial), and mitochondrial-read percentage (percent.mt). Red dashed lines indicate quality-control thresholds (nFeature_Spatial >50 and percent.mt <10%). The lower nFeature_Spatial threshold was set at 50 to retain biologically relevant regions with naturally low transcriptional activity, including terminally differentiated epithelial layers and HPV-associated lesions.

**Extended Data Fig. 6:**
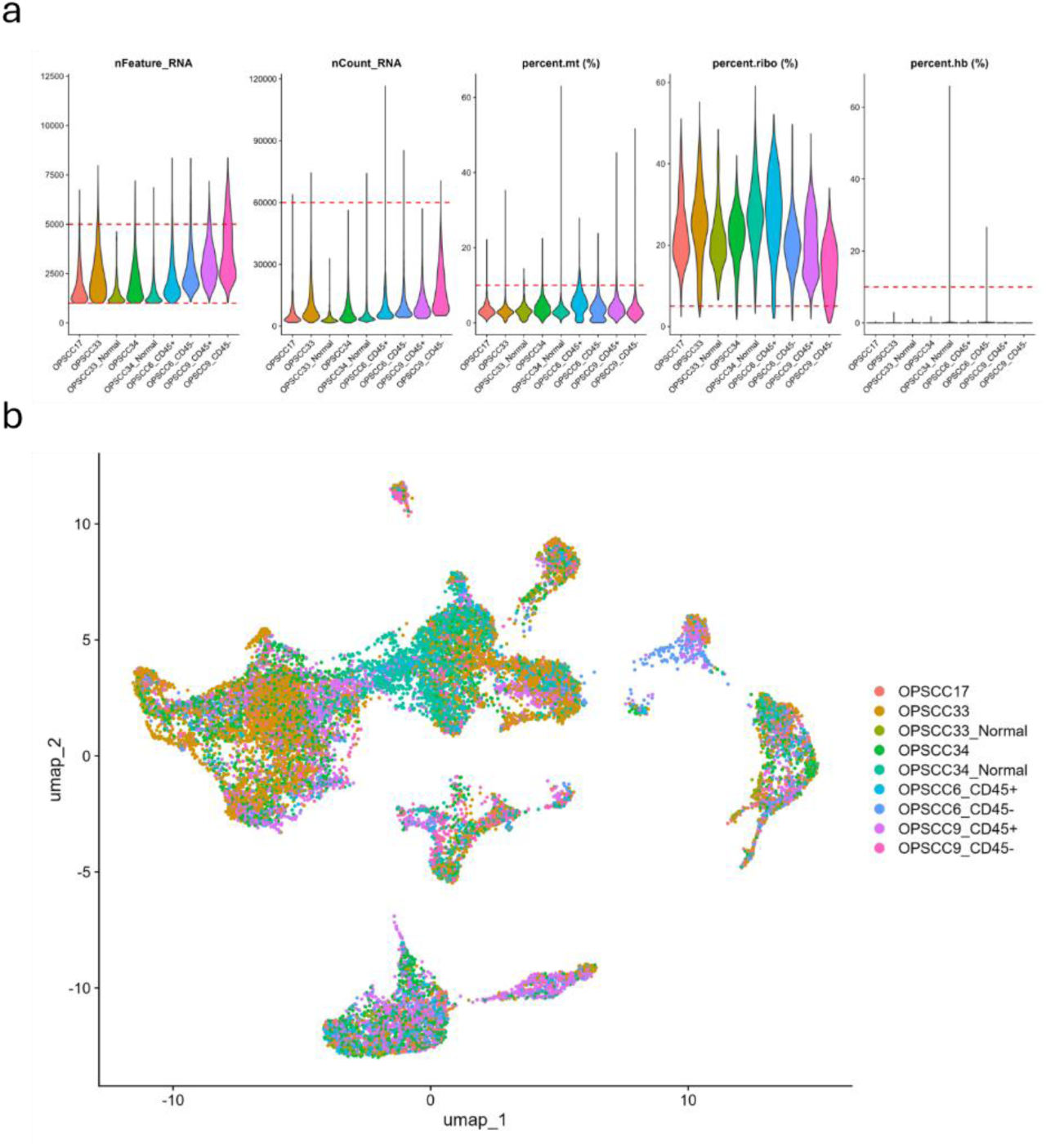
Quality control and integration of the single-cell RNA-seq dataset. **a.** Violin plots show quality-control metrics for the nine tonsillar single-cell libraries before downstream analysis. Samples are OPSCC17, OPSCC33 (Tumor and matched Normal), OPSCC34 (Tumor and matched Normal), and CD45-sorted fractions from OPSCC6 and OPSCC9 (CD45+ and CD45−). Metrics include detected genes (nFeature_RNA), total RNA counts (nCount_RNA), and percentages of reads mapping to mitochondrial (percent.mt), ribosomal (percent.ribo), and hemoglobin (percent.hb) genes. Horizontal red dashed lines indicate filtering thresholds used to exclude low-quality cells or artifacts. The lower nFeature_RNA threshold was 1,000, matching the prefiltering threshold of the original dataset. **b.** UMAP projection of all 20,995 filtered cells, color-coded by sample of origin as in a. Mixing of cells from different patients and tissue types (tumor versus normal) within clusters indicates successful reduction of batch effects during integration. OPSCC, oropharyngeal squamous cell carcinoma; UMAP, uniform manifold approximation and projection.

**Extended Data Fig. 7:**
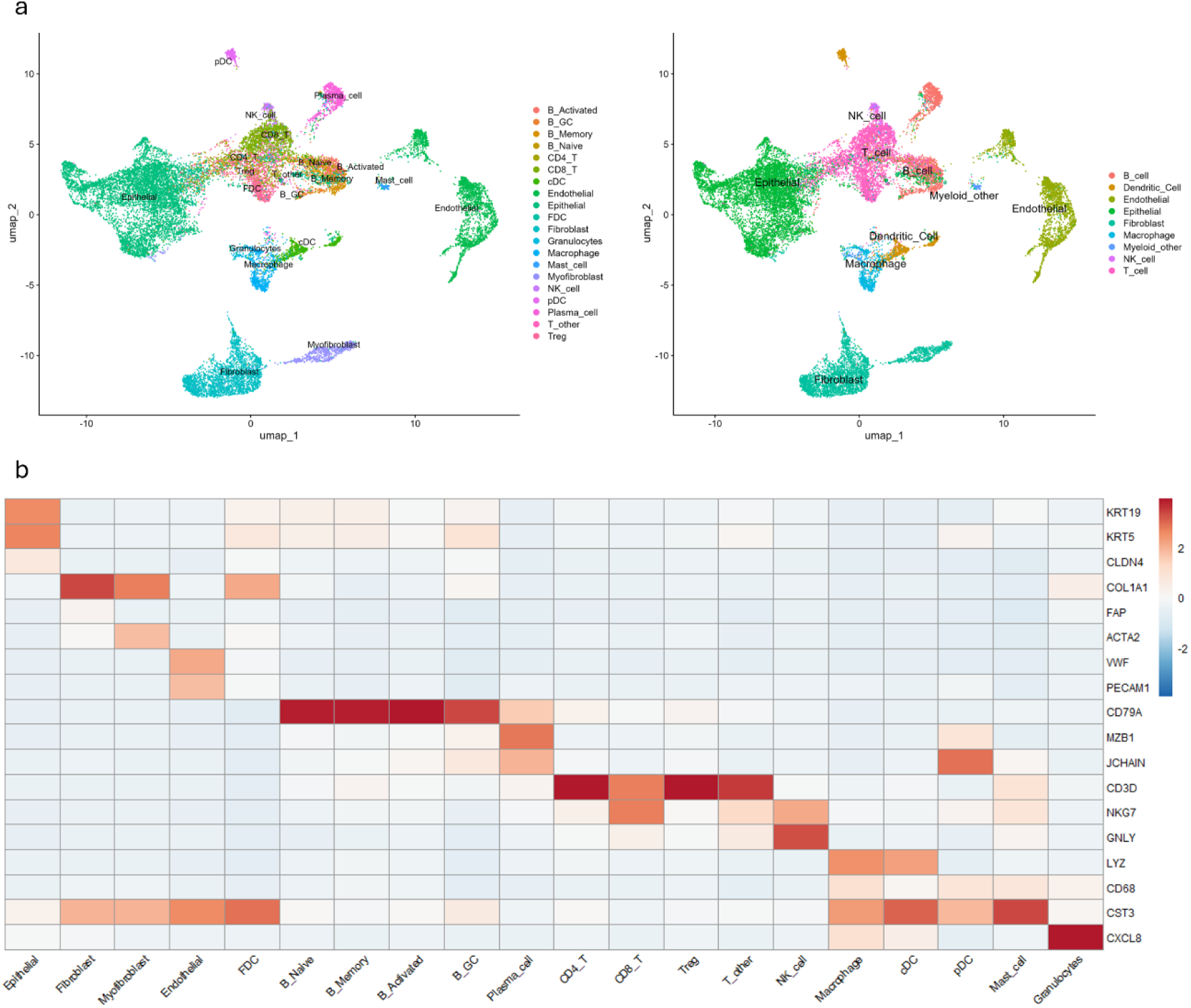
Cell-type annotation and marker gene validation of the scRNA-seq reference atlas. **a.** UMAP projections show hierarchical cell-type annotation of 20,995 high-quality cells from the integrated tonsillar dataset. Left, detailed cell states defined by a hybrid strategy: structural lineages (Epithelial, Fibroblast, Myofibroblast, Endothelial, and FDC) were manually annotated, whereas immune lineages (B-cell subsets, T-cell subsets, NK cells, and Myeloid lineages) were defined by Azimuth mapping. Right, broad lineages obtained by aggregating the detailed subclusters. **b.** Validation heatmap showing scaled average expression of canonical lineage markers across the final 20 high-resolution reference classes. Markers include KRT19/KRT5 (Epithelial), COL1A1/FAP (Fibroblast), ACTA2 (Myofibroblast), VWF/PECAM1 (Endothelial), CD79A (B cells), MZB1/JCHAIN (Plasma cells), CD3D (T cells), NKG7/GNLY (NK cells), LYZ/CD68/CST3 (Myeloid), and CXCL8 (Granulocytes). The strong diagonal pattern supports the specificity of the hybrid annotation workflow. cDC, classical dendritic cell; FDC, follicular dendritic cell; GC, germinal center; NK, natural killer; pDC, plasmacytoid dendritic cell; scRNA-seq, single-cell RNA sequencing; UMAP, uniform manifold approximation and projection.

**Extended Data Fig. 8:**
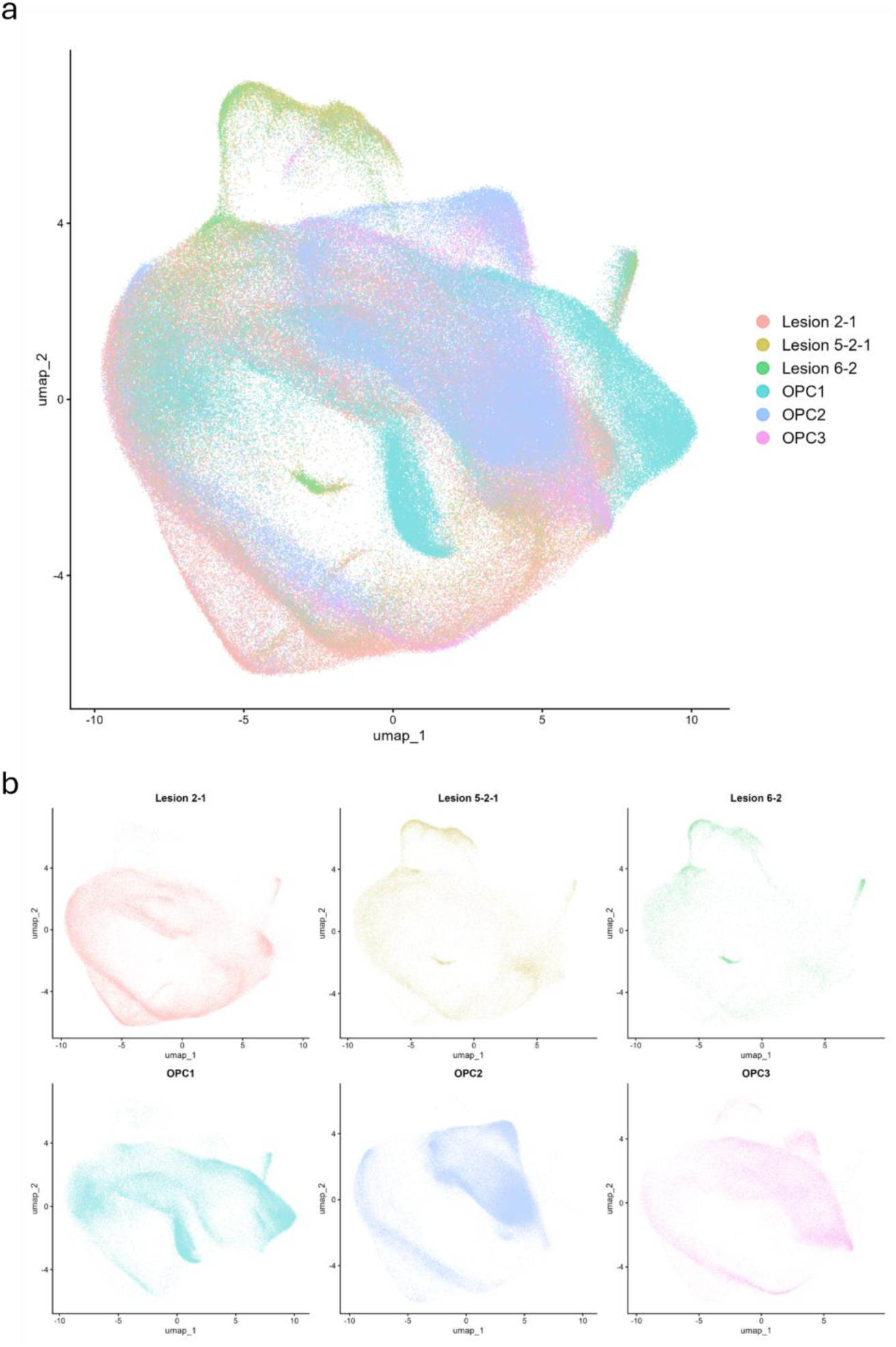
Integrated UMAP projection of high-purity epithelial bins. **a.** A total of 337,227 spatial bins from six cases with high epithelial purity (RCTD-estimated fraction >0.8) were integrated using Seurat’s RPCA workflow. Bins are color-coded by case of origin: intraepithelial lesions (Lesions 2-1, 5-2-1, and 6-2) and invasive tonsillar squamous cell carcinomas (OPC1, OPC2, and OPC3). Extensive overlap across the embedding supports successful reduction of batch and inter-individual effects. **b.** Case-specific UMAP projections showing each sample’s contribution to the integrated manifold. Axes and transcriptomic coordinates are identical to those in **a**. RCTD, robust cell type decomposition; RPCA, reciprocal principal component analysis; UMAP, uniform manifold approximation and projection.

**Extended Data Fig. 9:**
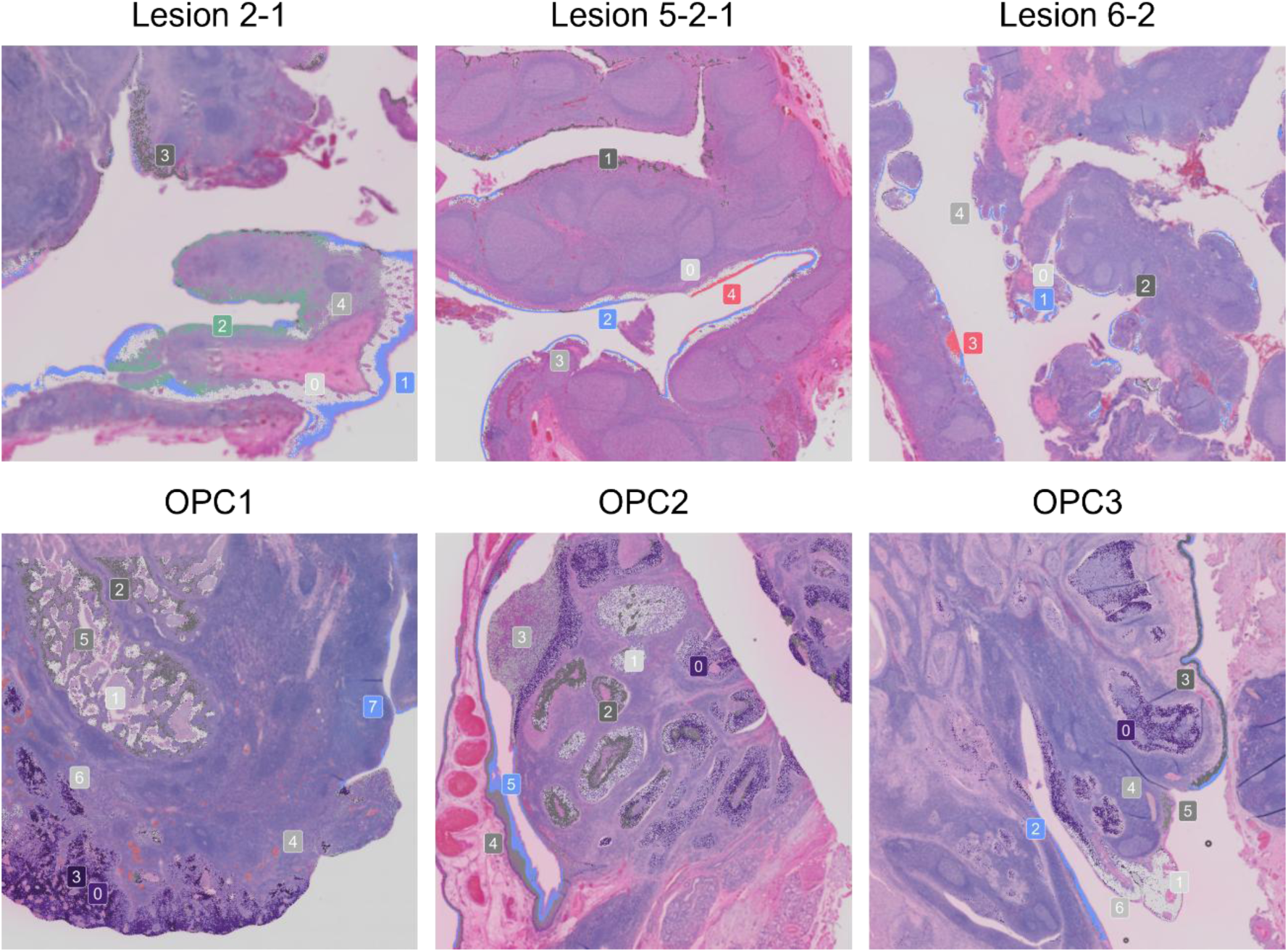
Spatial mapping of case-specific clusters and pathological annotations. Spatial transcriptomic clustering on HE sections. The top row shows the three lesions (Lesions 2-1, 5-2-1, and 6-2), and the bottom row the three invasive carcinoma cases (OPC1, OPC2, and OPC3). Each panel represents the 6.5 × 6.5 mm Visium HD capture area. Color-coded numbers overlaid on HE sections denote clusters identified independently within each sample; cluster numbers are case-specific and do not imply shared transcriptomic identities across samples. Pathological annotation: Based on expert review of HE morphology and spatially resolved transcriptomes, clusters within each case were assigned to four grades: Normal, Low-grade dysplasia, High-grade dysplasia, or Invasive Cancer. Summary of Pathological Grade Assignments: Normal: Lesion 2-1 (cluster 1), Lesion 5-2-1 (cluster 2), Lesion 6-2 (cluster 1), OPC1 (cluster 7), OPC2 (cluster 5), OPC3 (cluster 2) Low-grade Dysplasia: Lesion 5-2-1 (cluster 4), Lesion 6-2 (cluster 3) High-grade Dysplasia: Lesion 2-1 (cluster 2) Invasive Cancer: OPC1 (cluster 3), OPC2 (cluster 0), OPC3 (cluster 0)

**Extended Data Table 1.**
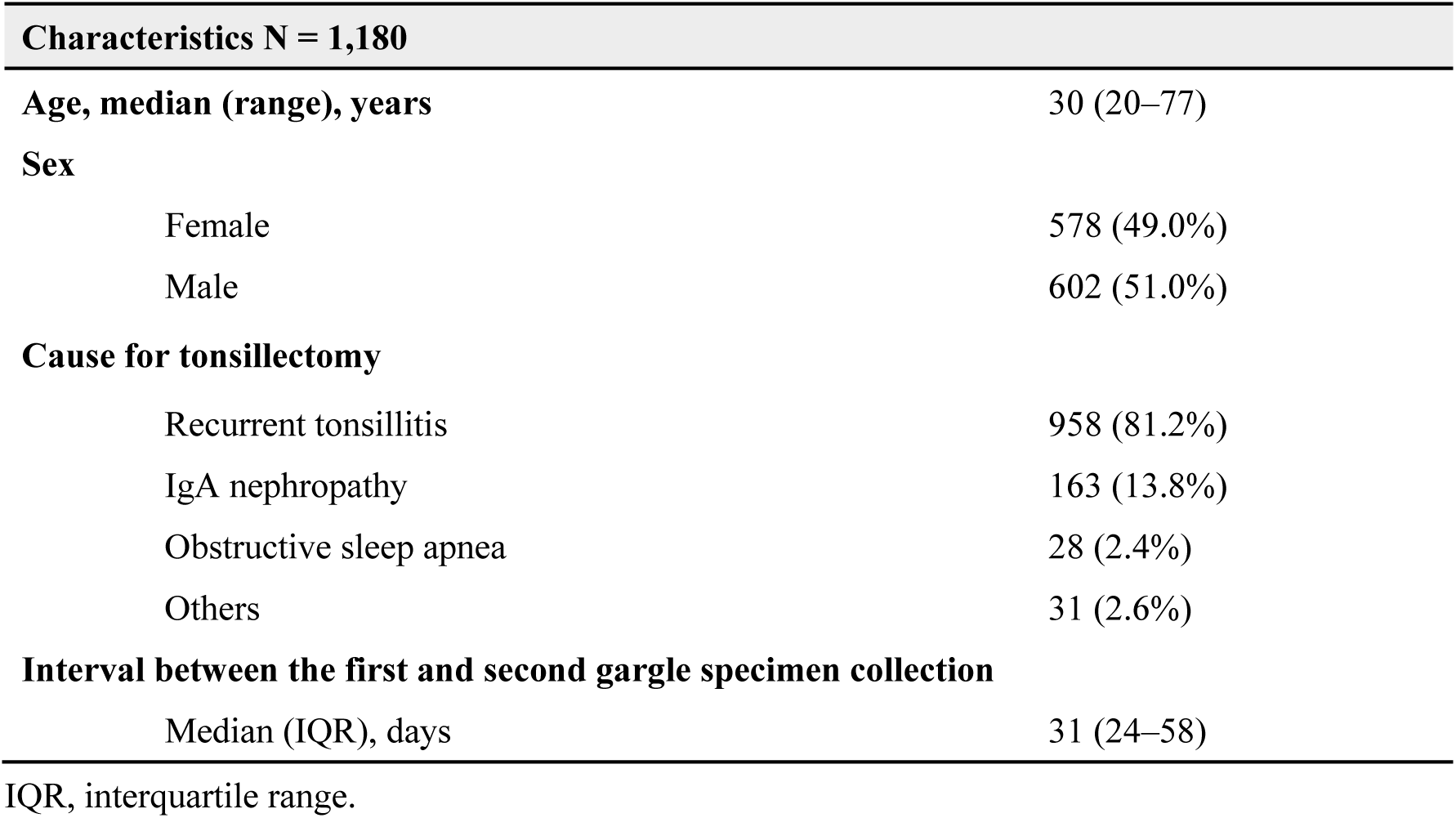
Baseline characteristics of patients.

| Characteristics N = 1,180 |  |  |
| --- | --- | --- |
| Age, median (range), years |  | 30 (20–77) |
| Sex |  |  |
| Female |  | 578 (49.0%) |
| Male |  | 602 (51.0%) |
| Cause for tonsillectomy |  |  |
| Recurrent tonsillitis |  | 958 (81.2%) |
| IgA nephropathy |  | 163 (13.8%) |
| Obstructive sleep apnea |  | 28 (2.4%) |
| Others |  | 31 (2.6%) |
| Interval between the first and second gargle specimen collection |  |  |
| Median (IQR), days |  | 31 (24–58) |

IQR, interquartile range.

**Extended Data Table 2.**
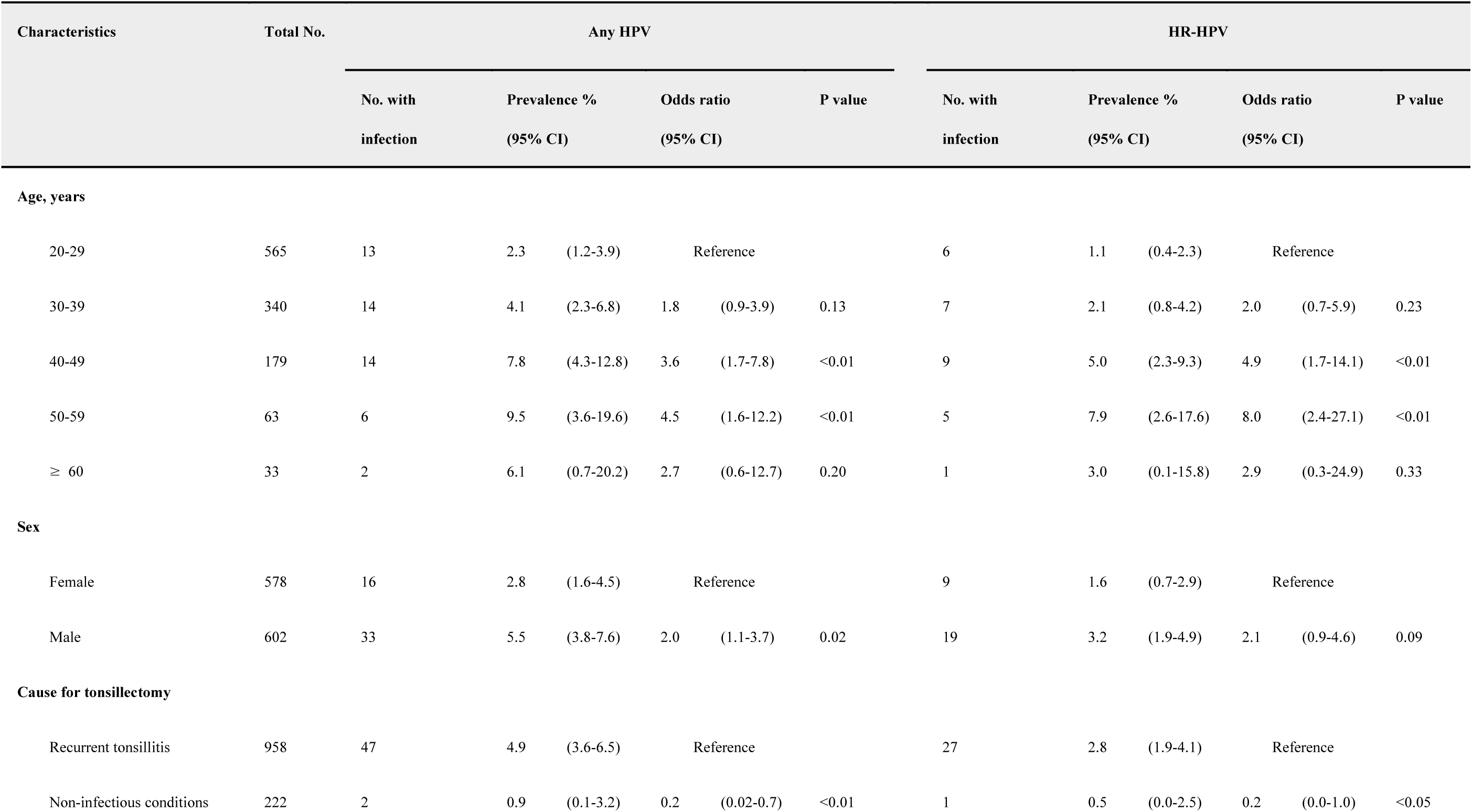
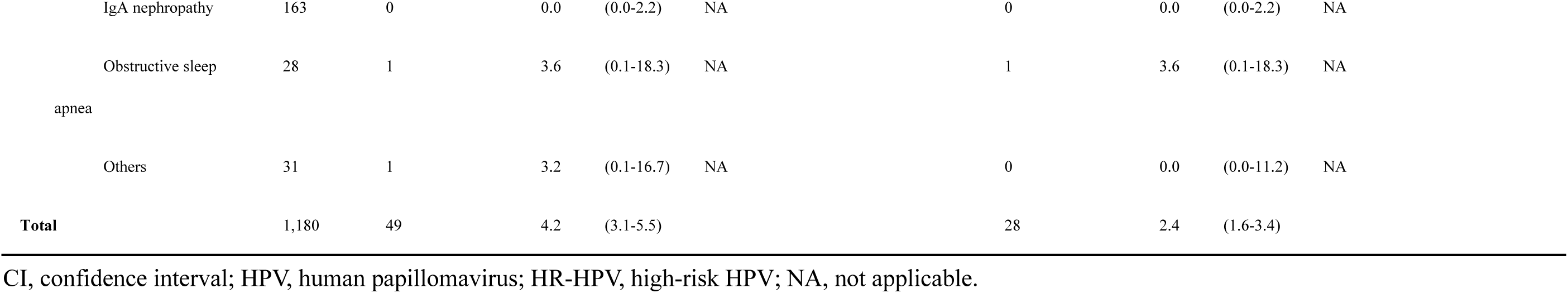
Prevalence of HPV infection according to the characteristics of patients.

**Extended Data Table 3.**
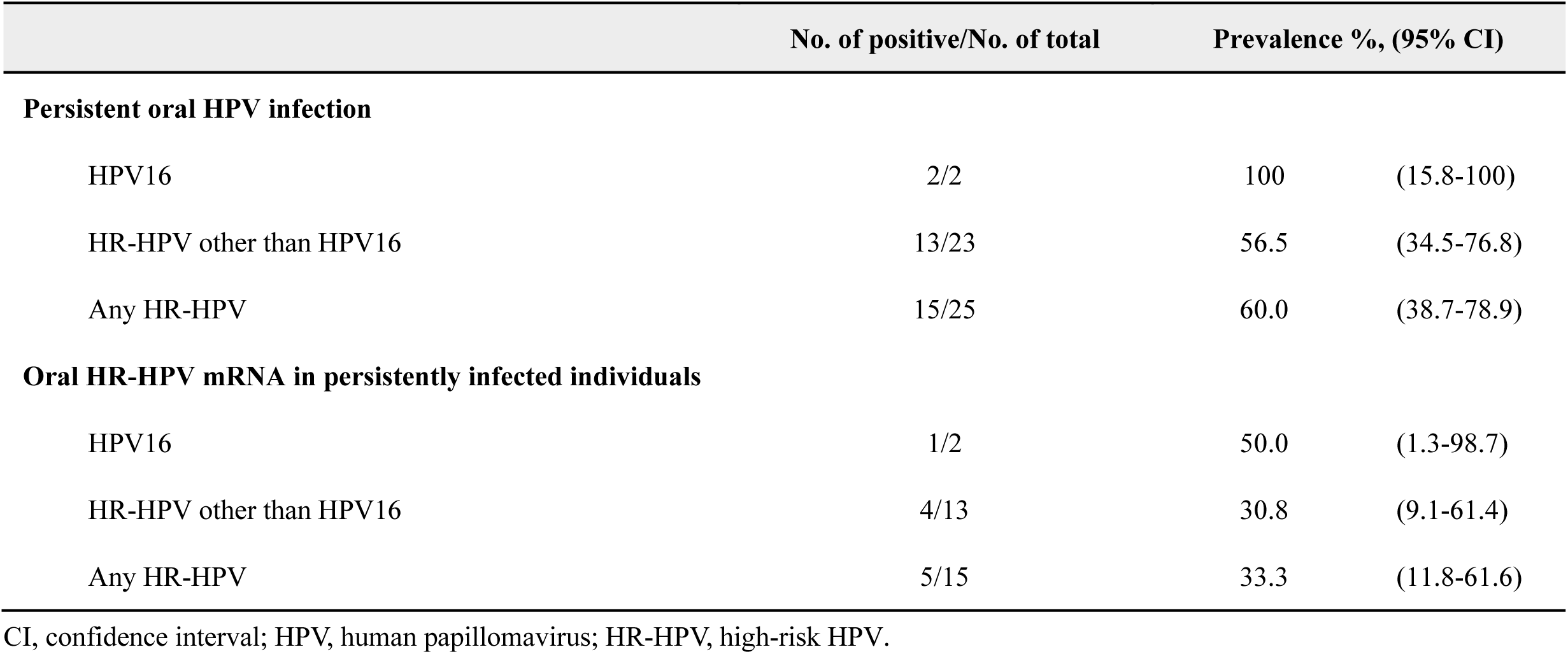
Persistent oral HPV infection and oral HPV mRNA status.

|  | No. of positive/No. of total | Prevalence %, (95% CI) |  |
| --- | --- | --- | --- |
| Persistent oral HPV infection |  |  |  |
| HPV16 | 2/2 | 100 | (15.8-100) |
| HR-HPV other than HPV16 | 13/23 | 56.5 | (34.5-76.8) |
| Any HR-HPV | 15/25 | 60.0 | (38.7-78.9) |
| Oral HR-HPV mRNA in persistently infected individuals |  |  |  |
| HPV16 | 1/2 | 50.0 | (1.3-98.7) |
| HR-HPV other than HPV16 | 4/13 | 30.8 | (9.1-61.4) |
| Any HR-HPV | 5/15 | 33.3 | (11.8-61.6) |

886 CI, confidence interval; HPV, human papillomavirus; HR-HPV, high-risk HPV.

**Extended Data Table 4.**
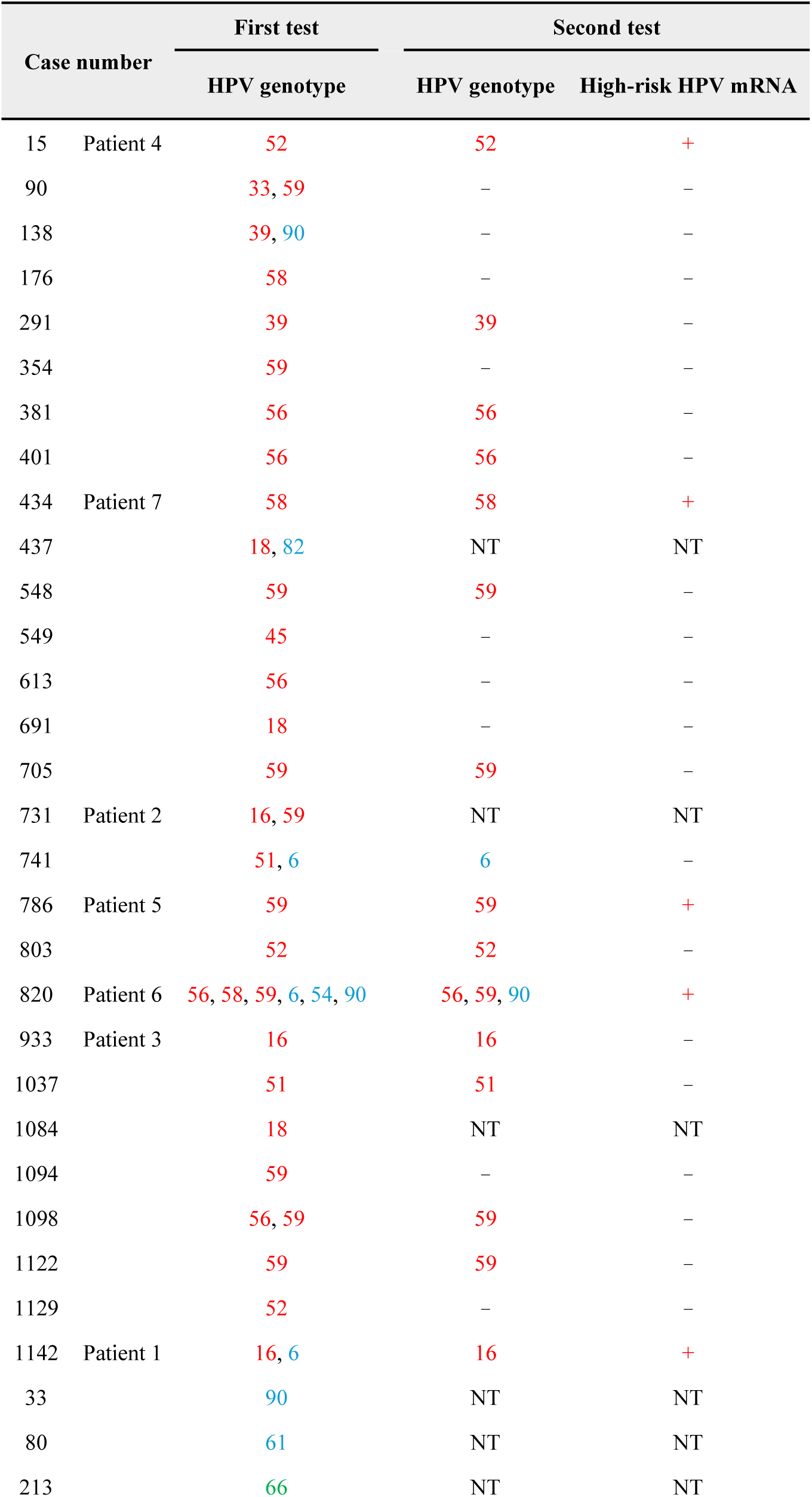
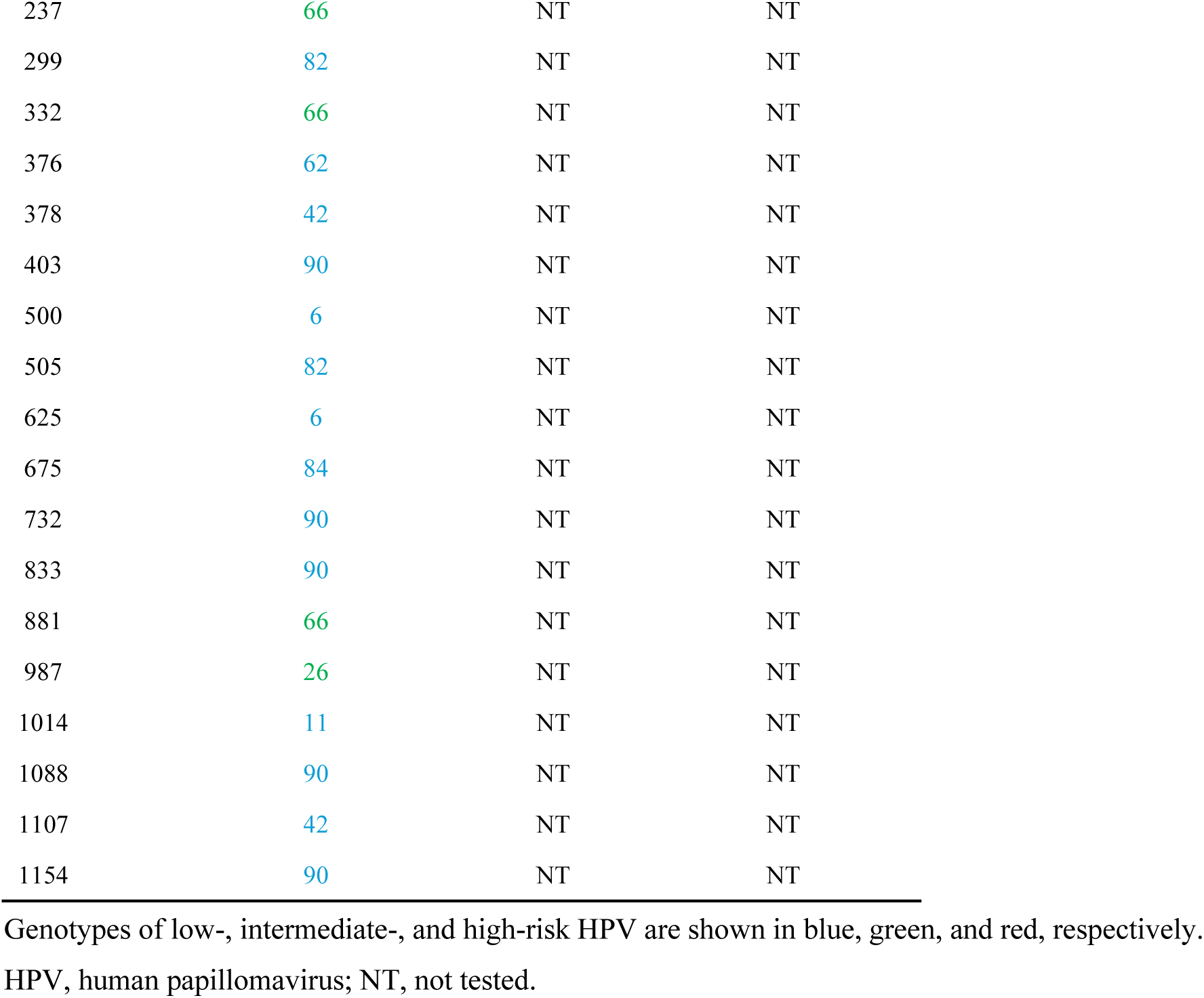
HPV genotype-specific profiles and high-risk HPV mRNA status in patients with oral HPV infection.

| Case number |  | First test | Second test |  |
| --- | --- | --- | --- | --- |
|  |  | HPV genotype | HPV genotype | High-risk HPV mRNA |
| 15 | Patient 4 | 52 | 52 | + |
| 90 |  | 33, 59 | – | – |
| 138 |  | 39, 90 | – | – |
| 176 |  | 58 | – | – |
| 291 |  | 39 | 39 | – |
| 354 |  | 59 | – | – |
| 381 |  | 56 | 56 | – |
| 401 |  | 56 | 56 | – |
| 434 | Patient 7 | 58 | 58 | + |
| 437 |  | 18, 82 | NT | NT |
| 548 |  | 59 | 59 | – |
| 549 |  | 45 | – | – |
| 613 |  | 56 | – | – |
| 691 |  | 18 | – | – |
| 705 |  | 59 | 59 | – |
| 731 | Patient 2 | 16, 59 | NT | NT |
| 741 |  | 51, 6 | 6 | – |
| 786 | Patient 5 | 59 | 59 | + |
| 803 |  | 52 | 52 | – |
| 820 | Patient 6 | 56, 58, 59, 6, 54, 90 | 56, 59, 90 | + |
| 933 | Patient 3 | 16 | 16 | – |
| 1037 |  | 51 | 51 | – |
| 1084 |  | 18 | NT | NT |
| 1094 |  | 59 | – | – |
| 1098 |  | 56, 59 | 59 | – |
| 1122 |  | 59 | 59 | – |
| 1129 |  | 52 | – | – |
| 1142 | Patient 1 | 16, 6 | 16 | + |
| 33 |  | 90 | NT | NT |
| 80 |  | 61 | NT | NT |
| 213 |  | 66 | NT | NT |

|  |  |  |  |
| --- | --- | --- | --- |
| 237 | 66 | NT | NT |
| 299 | 82 | NT | NT |
| 332 | 66 | NT | NT |
| 376 | 62 | NT | NT |
| 378 | 42 | NT | NT |
| 403 | 90 | NT | NT |
| 500 | 6 | NT | NT |
| 505 | 82 | NT | NT |
| 625 | 6 | NT | NT |
| 675 | 84 | NT | NT |
| 732 | 90 | NT | NT |
| 833 | 90 | NT | NT |
| 881 | 66 | NT | NT |
| 987 | 26 | NT | NT |
| 1014 | 11 | NT | NT |
| 1088 | 90 | NT | NT |
| 1107 | 42 | NT | NT |
| 1154 | 90 | NT | NT |

Genotypes of low-, intermediate-, and high-risk HPV are shown in blue, green, and red, respectively.
HPV, human papillomavirus; NT, not tested.

**Extended Data Table 5.** Clinicopathological characteristics of patients with oropharyngeal cancer.

| Case | Age/Sex | Subsite | Histology | p16 | HPV genotype | Stage* | Smoking history |
| --- | --- | --- | --- | --- | --- | --- | --- |
| OPC1 | 59/Male | tonsil | SCC | + | 16 | T2N1 | former (40 pack-years) |
| OPC2 | 57/Male | tonsil | SCC | + | 16 | T2N0 | never |
| OPC3 | 77/Female | tonsil | SCC | + | 16 | T2N1 | never |

\* According to the 8th edition of UICC TNM classification.
HPV, human papillomavirus; SCC, squamous cell carcinoma; UICC, Union for International Cancer Control.

**Extended Data Table 6.** Number and proportion of retained epithelial bins across different purity thresholds.

| Sample | Total bins | >50% |  | >60% |  | >70% |  | >80% |  | >90% |  |
| --- | --- | --- | --- | --- | --- | --- | --- | --- | --- | --- | --- |
|  |  | n (%) |  | n (%) |  | n (%) |  | n (%) |  | n (%) |  |
| Lesion 2-1 | 349283 | 114665 | (32.8) | 104675 | (30.0) | 90948 | (26.0) | 68872 | (19.7) | 32755 | (9.4) |
| Lesion 5-2-1 | 484734 | 51780 | (10.7) | 43180 | (8.9) | 35438 | (7.3) | 27549 | (5.7) | 17871 | (3.7) |
| Lesion 6-2 | 330029 | 37000 | (11.2) | 29513 | (8.9) | 22332 | (6.8) | 14893 | (4.5) | 6409 | (1.9) |
| OPC1 | 501495 | 179256 | (35.7) | 155358 | (31.0) | 124680 | (24.9) | 85316 | (17.0) | 37823 | (7.5) |
| OPC2 | 392171 | 197400 | (50.3) | 179258 | (45.7) | 148616 | (37.9) | 100934 | (25.7) | 44630 | (11.4) |
| OPC3 | 223794 | 73358 | (32.8) | 64883 | (29.0) | 53819 | (24.0) | 39663 | (17.7) | 21916 | (9.8) |

Total bins represent all spatial bins captured within each tissue section. Data are presented as the absolute number (n) and percentage (%) of bins meeting various estimated epithelial proportion thresholds (ranging from >50% to >90%) based on deconvolution analysis.

**Extended Data Table 7.**
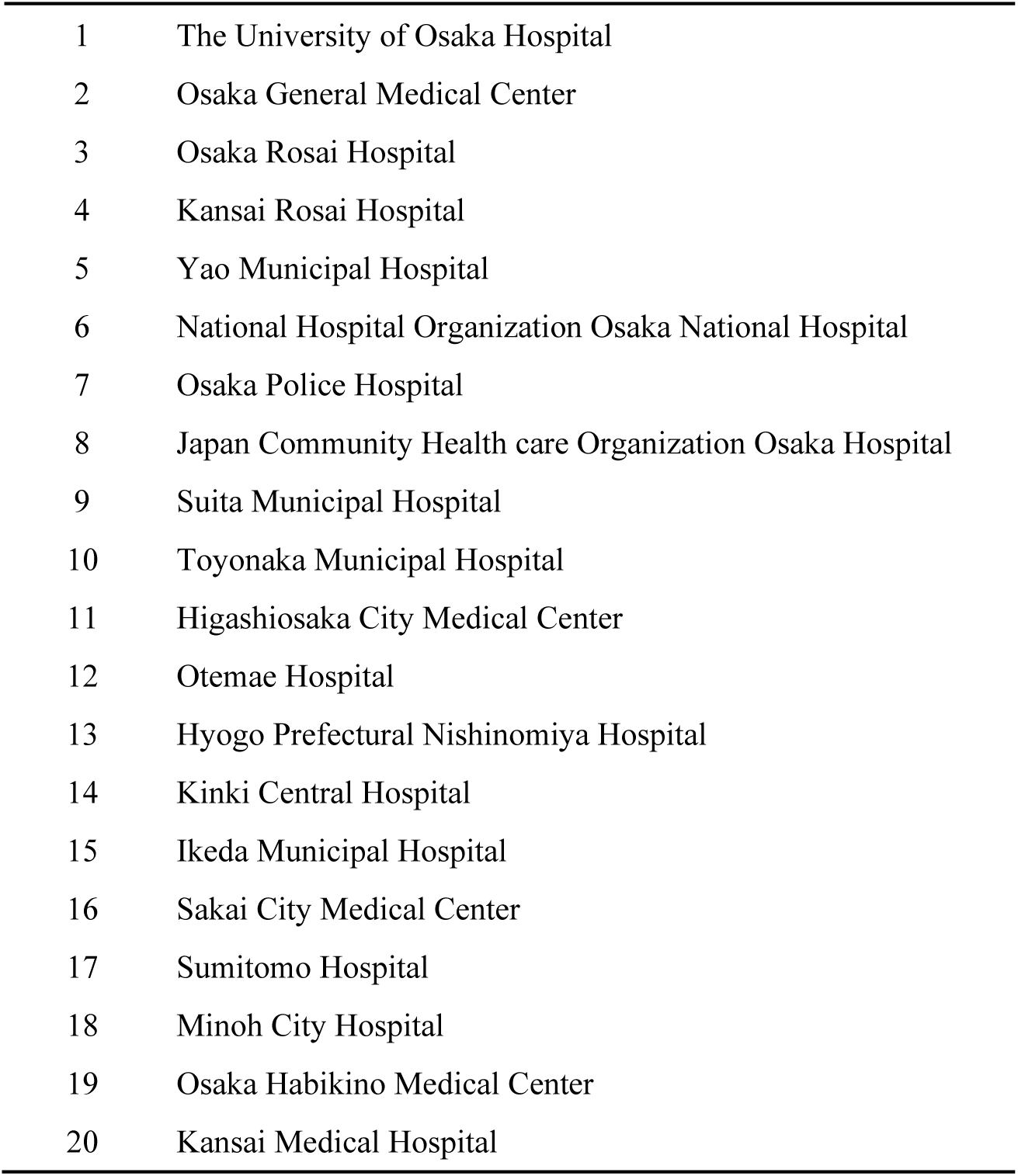
List of participating hospitals.

**Extended Data Table 8.**
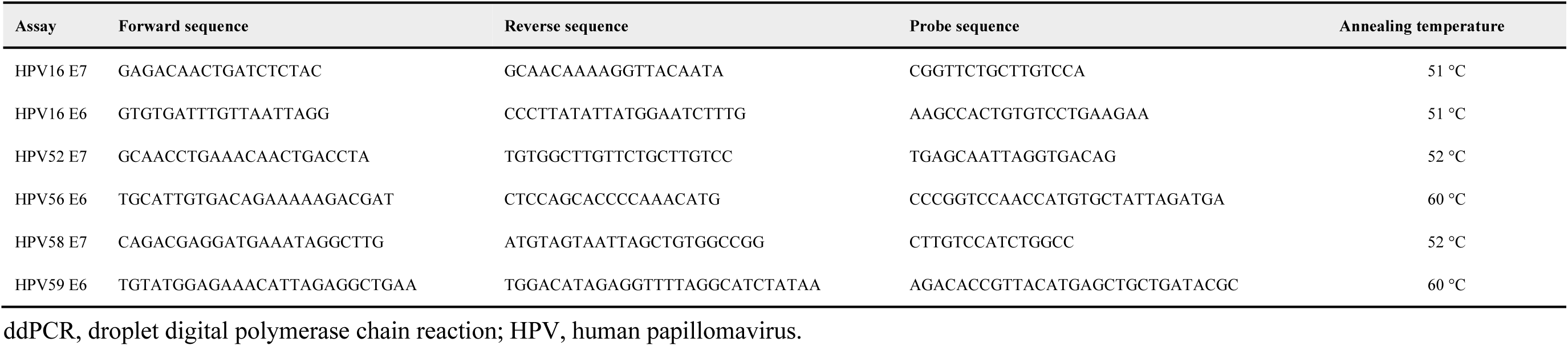
Oligonucleotide sequences used for ddPCR.

**Extended Data Table 9.**
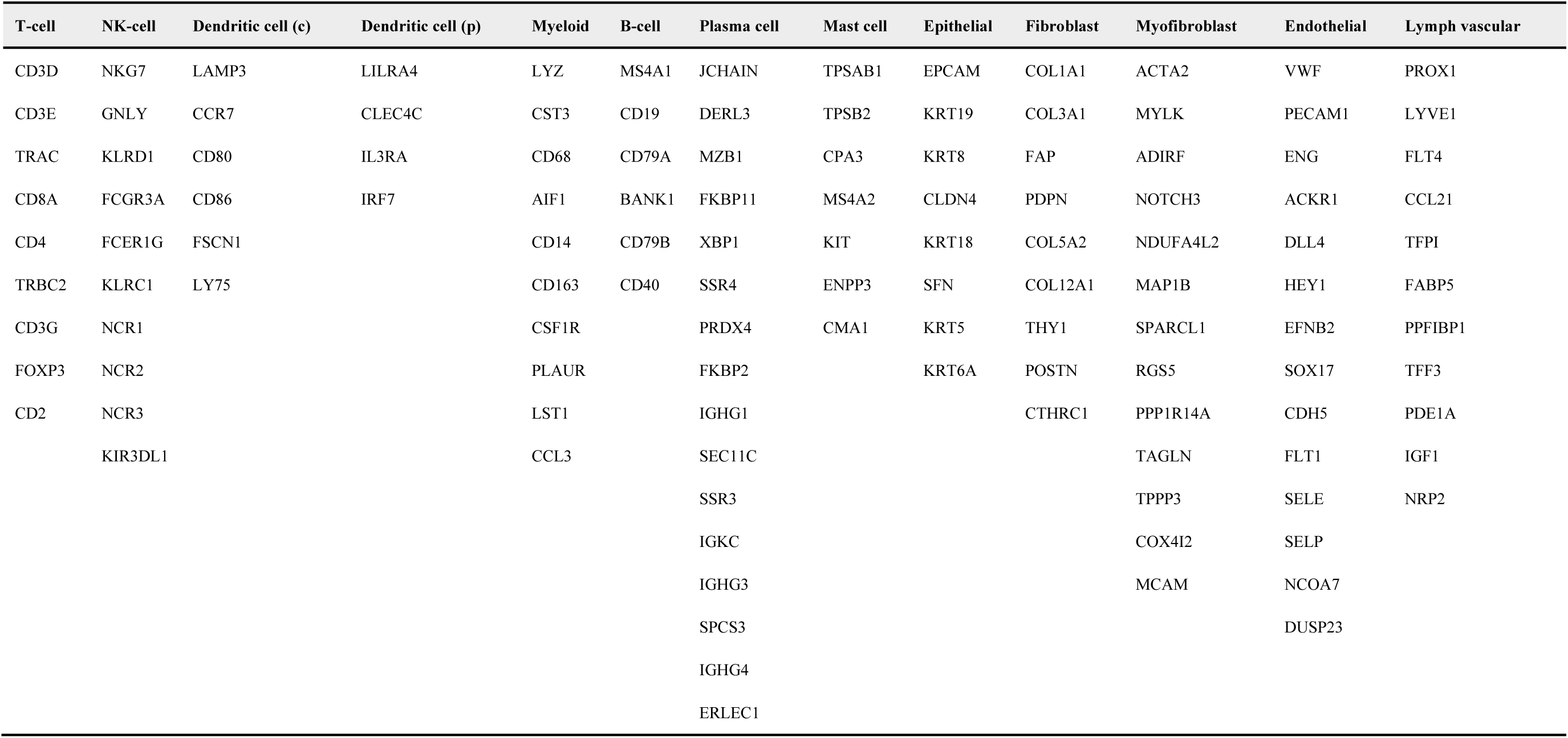
Marker genes of cell types and cellular states.

**Extended Data Table 10.** Gene signatures used for the calculation of the p-EMT Net Score.

| p-EMT Signature genes | Epithelial differentiation genes |
| --- | --- |
| LAMC2 | LYPD3 |
| PLAU | KRT16 |
| TNFRSF12A | SPRR1B |
| EMP3 | IL1RN |
| S100A2 | RHCG |
| LAMB3 | KRT6B |
| PTHLH | SLPI |
| TGFBI | CLDN7 |
| PHLDA1 | SDCBP2 |
| NDRG1 | THBD |
| LAMA3 | S100A8 |
| ITGA6 | CLDN4 |
| ITGA3 | PRSS8 |
| IGFBP7 | ELF3 |
| KRT17 | DSC2 |
| CDH3 | PVRL4 |
| P4HA2 | CDKN1A |
| TPM1 | GRHL1 |
| ARPC1B | S100A9 |
| PDPN | FABP5 |
| ACTN1 | DSG3 |
| CTSB | DMKN |

